# Stromal-cell deletion of STAT3 protects mice from kidney fibrosis by inhibiting pericytes trans-differentiation and migration

**DOI:** 10.1101/2021.08.19.456872

**Authors:** Amrendra K. Ajay, Li Zhao, Shruti Vig, Mai Fujikawa, Sudhir Thakurela, Shreyas Jadhav, I-Jen Chiu, Yan Ding, Krithika Ramachandran, Arushi Mithal, Aanal Bhatt, Pratyusha Chaluvadi, Manoj K. Gupta, Venkata S. Sabbisetti, Ana Maria Waaga-Gasser, Gopal Murugaiyan, David A. Frank, Joseph V. Bonventre, Li-Li Hsiao

## Abstract

Signal transducer and activator of transcription 3 (STAT3) is a key transcription factor implicated in the pathogenesis of kidney fibrosis. Although tubular *Stat3* deletion in tubular epithelial cells is known to protect mice from kidney fibrosis, the exact function of STAT3 in stromal cells remains unknown. We utilized stromal-cell *Stat3* knock-out (KO) mice, CRISPR and pharmacologic activators and inhibitors of STAT3 to investigate its function in pericyte-like cells. STAT3 is phosphorylated in tubular epithelial cells in acute kidney injury whereas its activation expanded to interstitial cells in chronic kidney disease in mice and humans. Stromal cell-specific deletion of *Stat3* protects mice from folic acid- and aristolochic acid-induced kidney fibrosis. Mechanistically, STAT3 directly regulates the inflammatory pathway, differentiation of pericytes into myofibroblasts. Specifically, STAT3 activation leads to an increase in migration and profibrotic signaling in genome-edited pericyte-like cells, 10T1/2. Conversely, *Stat3* KO or blocking STAT3 function inhibits detachment, spreading, migration, and profibrotic signaling. Furthermore, STAT3 binds to *Collagen1a1* promoter of fibrotic mouse kidneys and in pericyte-like cells. Together, our study identifies a previously unknown function of STAT3 in stromal cells that promotes kidney fibrosis and may have therapeutic value in fibrotic kidney disease.

**Graphical Abstract:** 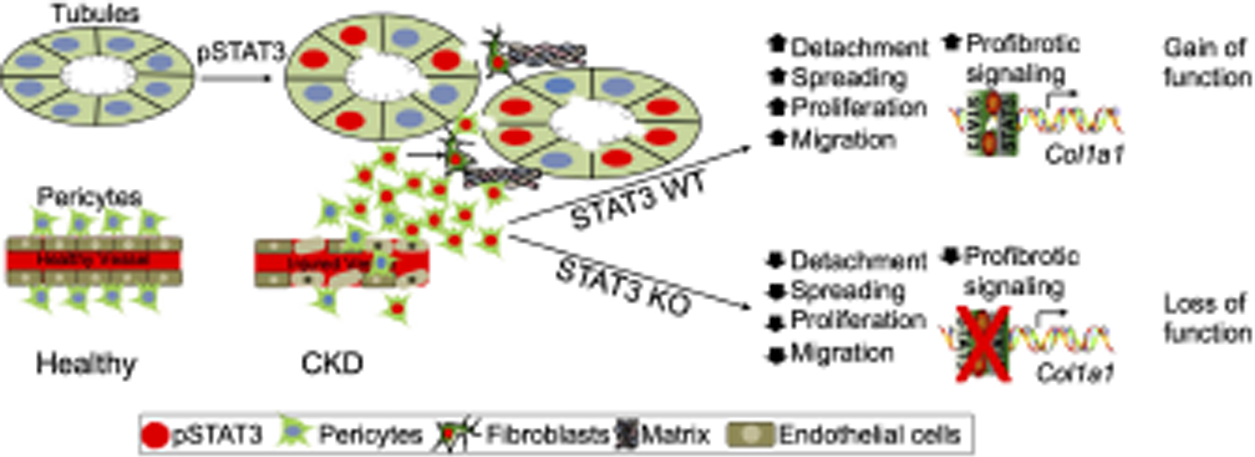

## INTRODUCTION

Diverse cell types play crucial functions in the cellular processes of tissue fibrosis, (1–4) including kidney fibrosis(5–8). Although many cell types in the tubulo-interstitium of the kidneys produce extracellular matrix (ECM), fibroblasts being the principal cells, the contribution of other cell types are not well understood. During the fibrotic process of kidneys, fibroblasts undergo transformation into myofibroblasts, which are identified as α-Smooth Muscle Actin ( α-SMA) positive cells. One of the central questions is to delineate the origin, activation, and regulation of these matrix-producing myofibroblasts (9–14). Many reports have suggested that activation of interstitial fibroblasts, differentiation of pericytes, phenotypic conversion of tubular epithelial cells and endothelial cells and recruitment of circulating fibrocytes are involved in this process (6, 15–19). Proliferation of interstitial cells and activation of myofibroblast are the two main phenotypical changes acquired by myofibroblasts to induce fibrosis (20), which is caused by inflammation and release of cytokines in the diseased kidneys (21, 22).

Previously, ECM-producing myofibroblasts were thought to be derived from resident fibroblasts after kidney injury (23). More recently, due to the availability of common markers for different cell types, there has been immense amount of effort towards identifying the “weight” of contribution of specific cell types to the fibrosis process and evidence that myofibroblasts can originate from many different cell types (10, 24) is starting to develop. Recent reports indicate that pericytes are a critical source of myofibroblasts in fibrotic kidneys (11, 16, 25, 26). Pericytes are a subset of the stromal cells that partially cover capillary walls, thereby stabilizing the endothelium. Following kidney injury, pericytes get detached from the endothelium, undergo proliferation and migration, and transform into myofibroblasts (16, 25).

Multiple mechanisms could account for the loss of peritubular capillaries in the fibrotic kidneys, one of them being the detachment of pericytes causing pericyte deficiency and destabilization of the endothelium (11). A recent study showed that perivascular CD73 positive cells attenuate inflammation and interstitial fibrosis in the kidney (27). Thus, investigating pericytes detachment and its associated cellular signaling may provide mechanistic insights into the initiation and progression of kidney fibrosis.

Secretion of inflammatory cytokines, (TGF-β (28, 29), CTGF (30), IL-1b, TNF-α (31)) including IL-6 (32), after cellular injury triggers inflammatory pathways in the kidney causing deterioration of kidney function (33). In the inflammatory cells, IL-6 binds to growth receptors to phosphorylate Signal Transduction and Activator of Transcription 3 (STAT3), resulting in increased inflammation (34, 35). We and others have shown that STAT3 regulates the expression of Kidney Injury Molecule-1 (KIM-1) following acute kidney injury (36, 37) and fibrinogen gamma chain in fibrotic kidneys (38). Furthermore, we found that IL-6 mediates STAT3 phosphorylation in human proximal tubular epithelial cells in acute kidney injury (AKI) (36) and IL-6-mediated STAT3 is observed in fibroblasts *in vitro* (38). Genetic depletion of STAT3 from tubular epithelial cells, using *KSP Cre,* protects mice from nephrectomy induced kidney fibrosis by inhibiting tubulo-interstitial crosstalk (39).

However, the role of STAT3 in the stromal cells particularly in pericytes in the kidney fibrosis remains unknown. Previous studies show that Foxd1 Cre driver can be used to investigate the function of pericytes (16, 27, 40–42). In the present study we found that *Stat3* KO mice in the stromal cells are protected from folic acid and aristolochic acid induced kidney fibrosis. Our gain of function and loss of function studies revealed that STAT3 directly regulates the inflammatory pathway, DNA damage, proliferation, migration, and differentiation of pericytes into myofibroblasts. Furthermore, we found that STAT3 binds to the promoter of *Collagen1a1* gene to execute the fibrotic process. Together, our study defines a previously unknown function by which STAT3 in stromal cells promotes kidney fibrosis.

## RESULTS

### Increased phosphorylation of STAT3 in AKI and CKD in mouse and human kidneys

We obtained human kidney specimens from the Brigham and Women’s Hospital pathology core from healthy individuals, those with Acute Tubular Necrosis (ATN) as AKI, and Diabetic Nephropathy (DN) as CKD. We found that phosphorylation of STAT3 is increased in AKI and CKD kidneys when compared to healthy kidney. In humans, pSTAT3 expression is limited to tubular epithelial cells of patients with ATN, while pSTAT3 was found in both tubular epithelial and the interstitial cells of the patients with CKD (**Figure 1A, B, C**). Of Note, the total number of tubular epithelial cells in both AKI and CKD were not significantly altered but the number of interstitial cells were significantly increased in CKD (**Supplemental Figure 1A, B**). Analogous to humans, the phosphorylation of STAT3 was also increased in mouse kidneys with FA or AA-induced injury. Specifically, we found increased pSTAT3 in tubular epithelial cells in AKI (2 days post FA or 4 days post AA treatments) and in interstitial cells in fibrotic kidneys (7 days post FA or AA treatment) when compared to those without FA treatment (**Figure 1D, E, F, H, and I**).

**Figure 1.**
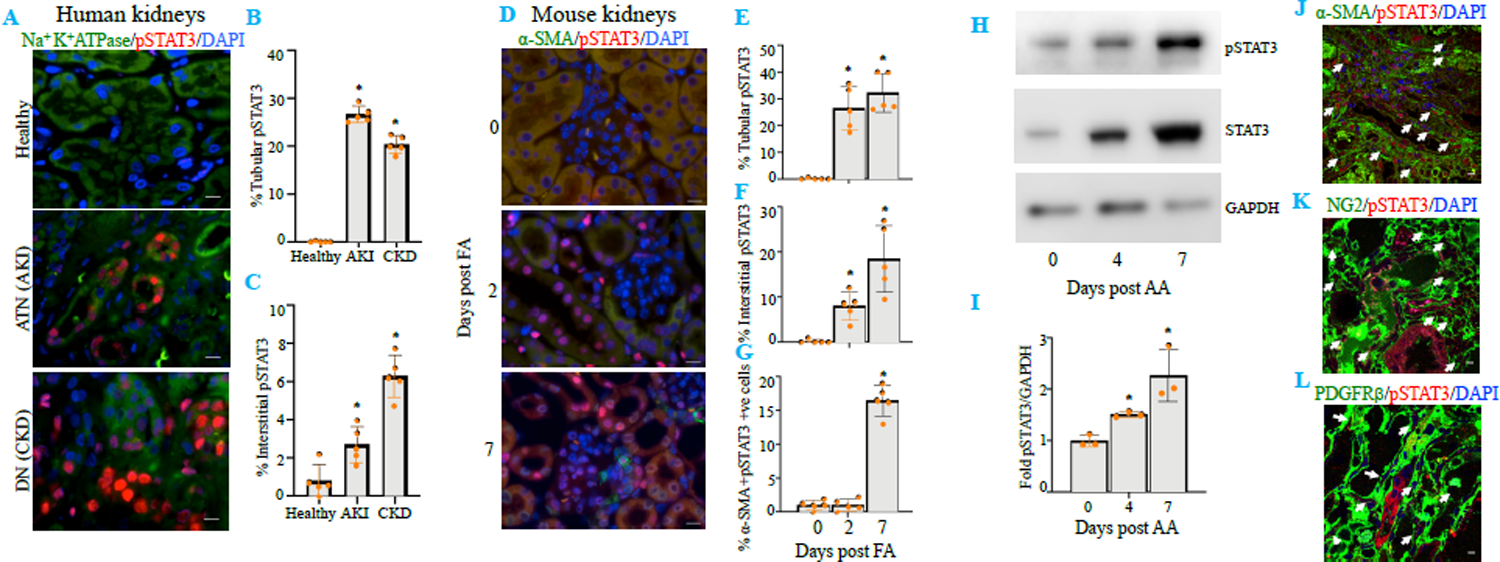
STAT3 is expressed in tubular epithelial cells in AKI and also in interstitial cells in CKD of human and mouse kidneys. (**A**) pSTAT3 staining (red) in healthy, AKI and CKD human kidneys. Na^+^K^+^ ATPase (green) for tubular staining. DAPI (blue) for nuclei. (**B**) Quantitation of number of pSTAT3 positive tubular epithelial cells. (**C**) Quantitation of number of pSTAT3 positive interstitial cells. * represents p<0.05 as compared to healthy. Scale bar=10 μm. (**D**) pSTAT3 (red) and α-SMA (green) in normal (day 0) and folic acid induced AKI (day 2) and fibrotic (day 14) kidneys. (**E**) Quantitation of percentage of tubular pSTAT3 positive cells. (**F**) Quantitation of percentage of interstitial pSTAT3 positive cells. (**G**) Quantitation of percentage of pSTAT3 and α-SMA positive cells. * represents p<0.05 as compared to normal. Scale bar=10 μm. (**H**) Western blotting for pSTAT3 and STAT3 in mouse normal (day 0) kidneys and aristolochic acid induced AKI (day 4) and fibrotic (day 7) kidneys. GAPDH served as loading control. (**I**) Quantitation of fold change in pSTAT3 compared to normal kidneys. * represents p<0.05 as compared to day 0. (**J, K, L**) Co-immunofluorescence staining for pSTAT3 and α-SMA or NG2 or PDGFRβ positive cells respectively in mouse kidneys 14 days post AA treatment. White arrow indicates double positive cells. Scale bar=10μm.

In addition, we found that interstitial myofibroblasts, which are known to play important roles in fibrosis, were positive for pSTAT3 as demonstrated by co-expression of a myofibroblast marker, α-SMA following 7 days post FA treatment (**Figure 1D and 1G**). Furthermore, we investigated the pSTAT3 positive cell types within the interstitium and found that in addition to myofibroblasts (**Figure 1J**), pericytes also have increased pSTAT3 as shown by co-immunostaining for either NG2 or PDGFRβ in AA induced fibrotic kidneys (**Figure 1K and 1L**) at 14 days post AA treatment.

### Stromal cell-specific deletion of *Stat3* protects mice from kidney fibrosis

A previous study showed that *Stat3* deletion from tubular epithelial cells protects mice from kidney fibrosis (39). We aimed to investigate the functions of STAT3 activation in the stromal cells, a cell population known to differentiate into myofibroblasts. To this aim, we developed a stromal cell specific *Stat3* deficient mice by crossing *Stat3* ^flox/flox^ mice and *Foxd1* ^GC^ mice (**Supplemental Figure 2A, 2B & 2C**). We did not find any significant difference in BUN and sCr in AKI (post FA treatment **Figure 2A, B** nor 4 days post AA treatment **Figure 2C, D**) between control (*Stat3 ^fl/fl^*) and *Stat3* KO (*Foxd1 ^GC^Stat3 ^fl/f^*^l^) mice. Furthermore, our results show similar tubular injury in the AKI phase for both control and *Stat3* KO mice (**Supplemental Figure 3A, D**); these observations were confirmed by mRNA levels of *Tnfa* and proximal tubular epithelial cell injury marker, *Hacvr1* suggesting that *Stat3* deletion in stromal cells did not affect the intensity of AKI (**Supplemental Figure 3B, C, E and F**).

**Figure 2.**
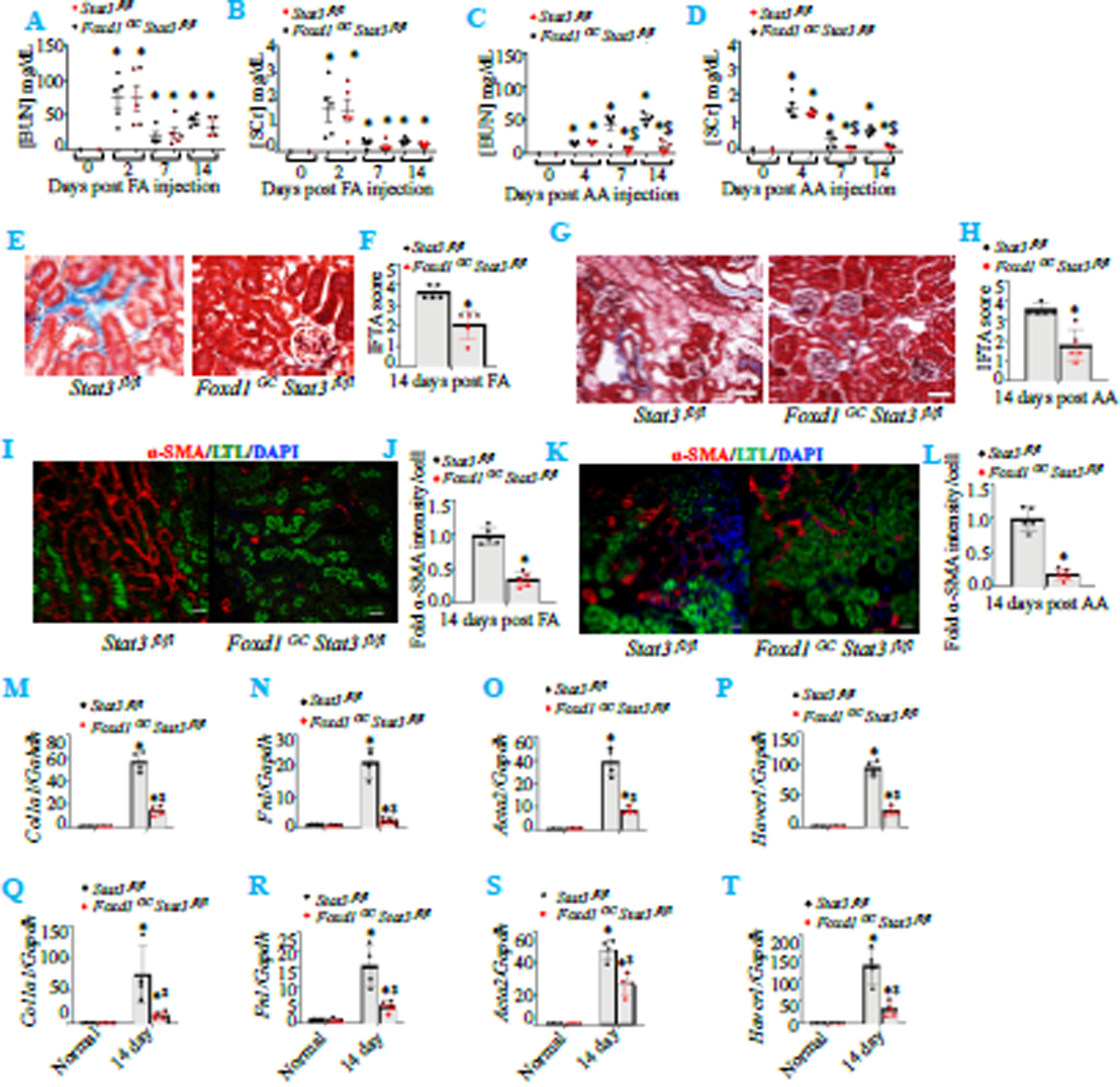
Stromal cell depletion of *Stat3* protects mice from folic acid and aristolochic acid induced kidney fibrosis. (**A-D**) BUN and SCr estimation for kidney dysfunction in FA-(**A, B**) or AA-(**C, D**) induced kidney injury. (**E-H**) Masson Trichrome Staining (MTS) and IFTA scoring of kidneys following 14 days post FA-(**E, F**) or AA-(**G, H**) induced kidney fibrosis. (**I-L**) Co-immunostaining for LTL (green) and α-SMA (red) and its quantitation in kidneys 14 days post FA-(**I, J**) or AA-(**K, L**) induced kidney fibrosis. (**M-T**) Gene expression analysis for *Col1a1*, *Fn1*, *Acta2* and *Havcr1* in FA-(**M, N, O** and **P**) or AA-(**Q, R, S** and **T**) induced kidney fibrosis. Scale bar=10 μm. * represents p<0.05 as compared to wildtype kidneys. ^$^ represents p<0.05 as compared to 14 days FA or AA wildtype kidneys.

We then assessed the functional roles of STAT3 in the fibrotic models using FA or AA treated mice for 7 days and 14 days. While our results revealed no significant difference in kidney function as assessed by BUN and sCr between control and *Stat3* KO mice in FA treated groups, we found that *Stat3* KO group had better kidney function in AA treated mice (**Figure 2A, B, C and D**). We then evaluated the extent of kidney injury by assessing the degree of fibrosis, myofibroblast activation and tubular damage in both FA and AA treatment groups. Using MTS staining, we found a significant decrease in fibrosis in *Stat3* KO mice as compared to control mice (**Figure 2E, F, G and H**). Myofibroblast activation, assessed by α-SMA staining, was also decreased in *Stat3* KO mice (**Figure 2I, J, K and L**). We further assessed the tubular function, by brush border using LTL staining, and found an increase in LTL staining in *Stat3* KO mice, indicating the preservation of tubular function (**Figure 2I, K**). In addition, we found a decrease in the expression of *Collagen1a1, Fibronectin1* and *Acta2* in *Stat3* KO mice when compared with control mice (**Figure 2M, N, O, Q, R and S**) in both FA- and AA-induced fibrosis. Furthermore, we found decreased gene expression of *Havcr1* in *Stat3* KO mice as compared to control mice (**Figure 2P, T**). Together, our results suggest that *Stat3* depletion from stromal cells has protective effects on kidney fibrosis in FA and AA treated models.

### Stromal cell-specific *Stat3* deletion results in reduced inflammation, DNA damage and cell proliferation in the fibrotic kidneys, *in vivo*

To understand the mechanisms by which stromal cell-specific deletion of *Stat3* protects mice from kidney fibrosis, we investigated the involvement of inflammatory pathways in the pericytes and macrophages in AA induced kidney fibrosis model. We found that infiltration of macrophages (stained by F4/80) in the kidney was significantly reduced in *Stat3* KO as compared to control mice (**Figure 3A, B**). We then isolated the macrophages from control and *Stat3* KO fibrotic mice kidneys using F4/80 and Cd11b markers and found decreased number of macrophages in *Stat3* KO mice kidneys as compared to control kidneys (**Figure 3C, D**). Furthermore, gene expression of inflammatory mediators including *Il1b*, *Il6*, *Il12*, *Il23*, *Ifng*, *Nos2* and *Tnfa,* was decreased in the macrophages of *Stat3* KO mice as compared to control mice (**Figure 3E**). On the other hand, expression levels of *Il10* and *Tgfb*, markers of M2 macrophages, a reparative phenotype, were increased in the fibrotic kidneys of *Stat3* KO mice (**Figure 3F**).

**Figure 3.**
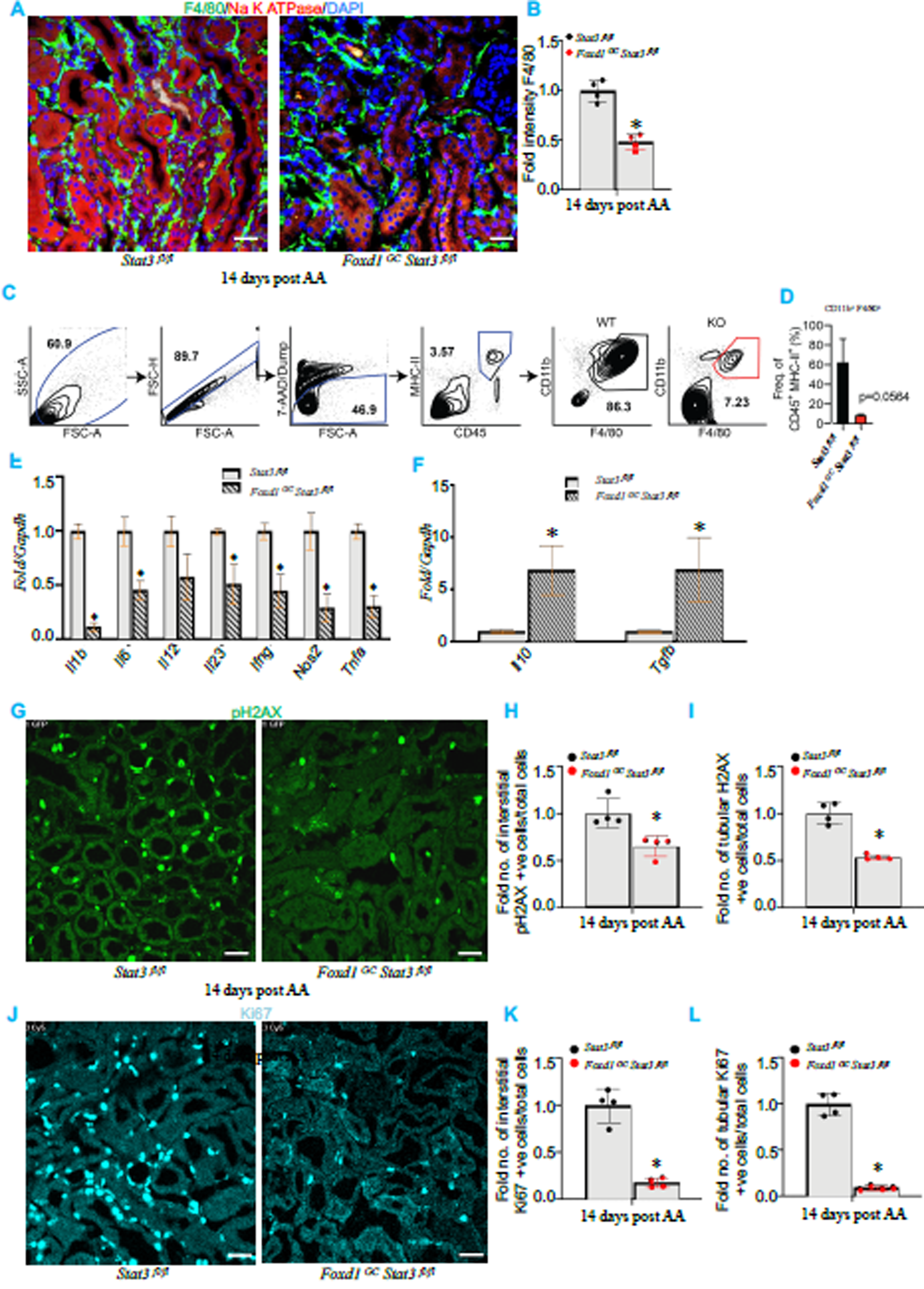
Stromal cell depletion of *Stat3* shows decreased macrophage infiltration, inflammation, decreased DNA damage signaling and reduced cell proliferation in fibrotic kidneys. (**A**) Co-immunostaining for F4/80 (green) and Na^+^ K^+^ ATPase in the mouse kidneys 14 days post AA injection. (**B**) Quantitation of F4/80 intensity. Scale bar=10 μm. (**C** and **D**) FACS sorting strategy for macrophages from fibrotic kidneys. Quantitative RT-PCR for (**E**) inflammatory mediators and (**F**) M2 markers from the macrophages isolated from kidney 14 days post AA-induced fibrosis. (**G**) Immunostaining for pH2AX (green) in the mouse kidneys 14 days post AA injection. Quantitation of (**H**) interstitial and (**I**) tubular pH2AX positive cells. (**J**) Immunostaining for Ki67 (Cyan) in the mouse kidneys 14 days post AA injection. Quantitation of (**K**) interstitial and (**L**) tubular Ki67 positive cells. Scale bar=10 μm. * represents p<0.05 as compared to wildtype kidneys.

Furthermore, to assess the involvement of STAT3 in DNA damage signaling as well as proliferation of tubular and interstitial cells, we stained for pH2AX and Ki67, respectively. We found that *Stat3* KO mice show reduced DNA damage of interstitial as well as tubular epithelial cells (**Figure 3G, H and I**). We also found decreased numbers of interstitial and tubular Ki67 positive cells in *Stat3* KO mice (**Figure 3J, K and L**). Taken together, our results indicated that the depletion of *Stat3* from stromal cells protects mice from kidney fibrosis by reducing inflammation, DNA damage, proliferation of the interstitial cells and increasing the polarization of M2 macrophages.

### IL-6 mediated STAT3 phosphorylation leads to increased proliferation, migration and profibrotic signaling in pericyte-like 10T1/2 cells, *in vitro*

It has been shown that stromal cells including pericytes can differentiate into myofibroblasts (16, 43–46). Due to the lack of a pericytes cell line, we utilized 10T1/2 cells of mesenchymal stem cell origin, which are widely used for studying pericyte biology in various organs including kidney (47–49). As an increase in IL-6 has been reported in the pathogenesis of kidney disease, (50–52) we explored its function in STAT3 mediated proliferation, migration, and induction of fibrotic signaling in 10T1/2 cells. We found that pSTAT3 is increased and translocated to the nucleus after IL-6 treatment, and this was inhibited by stattic, a specific inhibitor of STAT3, indicating that these cells responded to IL-6 (**Figure 4A, B, C and D**). We also found increased proliferation of 10T1/2 cells after IL-6 induced STAT3 activation, which was also inhibited by stattic as shown by BrdU incorporation and Ki67 staining (**Figure 4E, F, G and H**). Migration of pericytes to the site of injury is one of the critical steps for the development of fibrosis (12, 25). We found that an increase in the migration of 10T1/2 cells was associated with the phosphorylation of STAT3, as shown by scratch assay and trans well migration assay, which can be inhibited by stattic (**Figure 4I, J, K and L; Supplemental Figure 4**). We also report here that the phosphorylation of STAT3 increased the phosphorylation and nuclear translocation of profibrotic transcription factor SMAD2, which was also inhibited by stattic (**Figure 4M, N**). In addition, increased STAT3 phosphorylation by IL6 resulted in higher levels of the fibrotic markers, Collagen1, Fibronectin and α-SMA (**Figure 4O, P, Q, R, S and T**). Interestingly, static effectively inhibited IL-6-induced upregulation of these fibrotic markers. Thus, these data indicate that IL-6 mediated-STAT3 phosphorylation increases proliferation, migration and profibrotic signaling in 10T1/2 cells.

**Figure 4.**
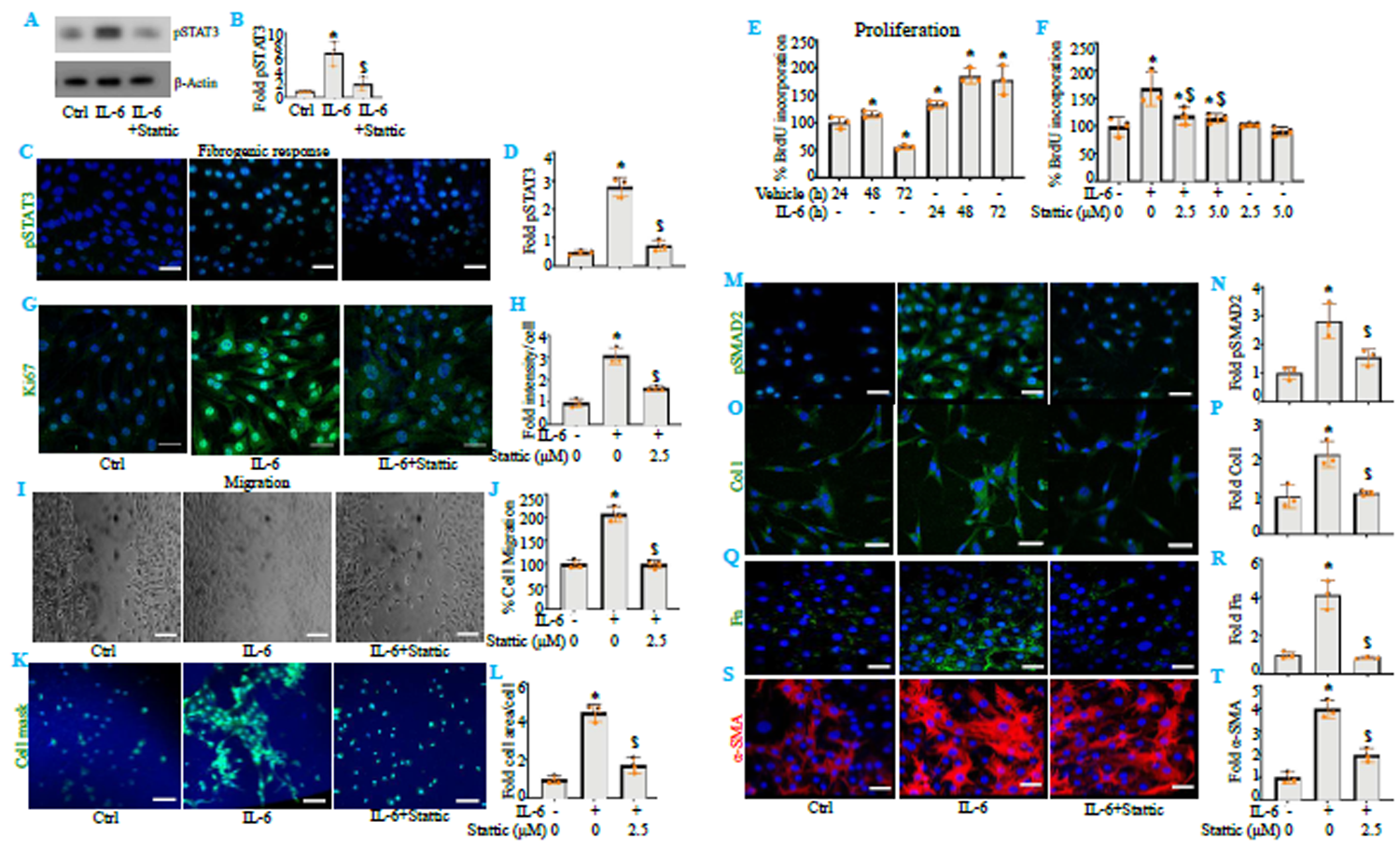
IL-6 mediated STAT3 phosphorylation regulates proliferation, migration and profibrotic signaling in pericyte-like 10T1/2 cells. (**A** and **B**) Western blotting for STAT3 phosphorylation following treatment with IL-6 or IL-6 in combination with stattic in 10T1/2 cells and its quantitation. (**C** and **D**) Nuclear translocation of STAT3 in 10T1/2 cells following IL-6 and or stattic treatment and its quantitation. (**E** and **F**) Proliferation of 10T1/2 cells following IL-6 and stattic treatments using BrdU assay in a (**E**) time and (**F**) STAT3 dependent manner. (**G** and **H**) Immunofluorescence staining for Ki67 with or without IL-6 and stattic treatments and its quantitation. (**I** and **J**) Migration of 10T1/2 cells following IL-6 and stattic treatment using scratch assay and its quantitation. (**K** and **L**) Trans well migration of 10T1/2 cells treated with IL-6 and stattic and its quantitation for cell area normalized to cell number. (**M** and **N**) Immunostaining for pSMAD2 (green) and its quantitation. (**O** and **P**) Immunostaining for Collagen 1 (Col1, green) and its quantitation. (**Q** and **R**) Immunostaining for Fibronectin (Fn, green) and its quantitation. (**S** and **T**) Immunostaining for α-SMA (green) and its quantitation. Scale bar=10 μm. * represents p<0.05 as compared to control cells. ^$^ represents p<0.05 as compared to IL-6 treated cells.

### STAT3 directly regulates proliferation, migration and profibrotic signaling in pericytes, *in vitro*

To investigate the function of STAT3 phosphorylation in pericytes, we established cellular models for gain and loss of STAT3 function. We generated CRISPR mediated STAT3 Synergistic Activation Mediator (SAM), that transcriptionally activates STAT3 expression as well as STAT3-C by mutating Alanine 662 and Asparagine 664 to Cystines that leads to dimerization and constitutive activation of STAT3 (**Figure 5A**). As expected, *Stat3* mRNA was significantly increased in SAM STAT3 with no change in STAT3-C (**Supplemental Figure 5**). SAM STAT3 and STAT3-C cells showed abundance of pSTAT3 as compared to control cells (**Figure 5B, C**). Furthermore, we found that the expression of profibrotic transcription factor pSMAD2 and fibrotic markers Collagen1, Fibronectin and α-SMA was increased in STAT3 activated cells as compared to control cells (**Figure 5D, E, F, G, H, I, J and K**). In addition, we tested the function of STAT3 in migration of 10T1/2 cells. STAT3 activated cells (SAM STAT3 and STAT3-C) showed increased trans well migration with increased cell area as compared to control cells (**Figure 5L, M and N**).

**Figure 5.**
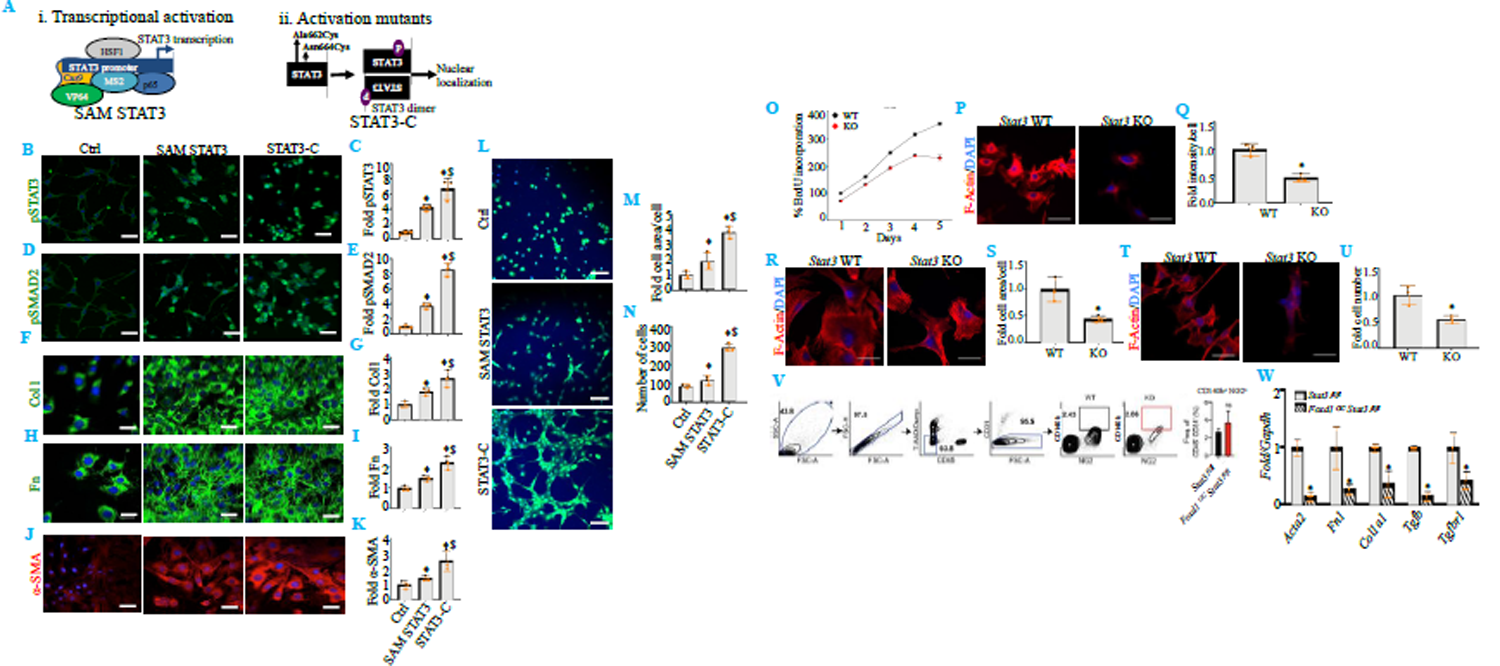
STAT3 directly regulates attachment, spreading, migration, proliferation and profibrotic signaling in 10T1/2 cells. (**A**) Scheme for STAT3 activation using synergistic activation mediators (SAM) and activation mutants (STAT3-C). (**B** and **C**) Immunostaining for pSTAT3 (green) and its quantitation. (**D** and **E**) Immunostaining for pSMAD2 (green) and its quantitation. (**F** and **G**) Immunostaining for Collagen 1 (Col1, green) and its quantitation. (**H** and **I**) Immunostaining for Fibronectin (Fn, green) and its quantitation. (**J** and **K**) Immunostaining for α-SMA (green) and its quantitation. (**L**) Trans well migration of 10T1/2 cells with STAT3 activation. Quantitation of (**M**) cell area normalized to cell number and (**N**) total number of migrated cells. Scale bar=10 μm. * Represents p<0.05 as compared to control cells. (**O**) Proliferation of *Stat3* WT and KO 10T1/2 cells as assayed by BrdU incorporation assay. (**P** and **Q**) Adherence assay (F-actin, red) and quantitation of intensity of cells normalized to total number of cells. (**R** and **S**) Spreading assay (F-actin, red) and quantitation of cell area normalized to total number of cells. (**T** and **U**) Trans well migration assay (F-actin, red) and quantitation of total number of migrated cells. (**V**) FACS sorting strategy for isolating pericytes from fibrotic mice kidneys and its quantitation. (**W**) Quantitative RT-PCR for pro-fibrotic factors in the pericytes isolated from fibrotic control and *Stat3* KO mice. Scale bar=10 μm. *represents p<0.05 as compared to wildtype 10T1/2 cells or control mice pericytes.

We then investigated whether STAT3 played direct roles in cell proliferation, cell detachment, cell spreading and cell migration. To investigate these phenotypes, we knocked out *Stat3* in 10T1/2 using CRISPR as a loss of function model. *Stat3* KO cells showed genomic deletion and absence of STAT3 protein as compared to *Stat3* WT cells (**Supplemental Figure 6**). Our results demonstrated that *Stat3* KO cells exhibited decreased proliferation, surface attachment, spreading and migration as compared to *Stat3* WT cells (**Figure 5O, P, Q, R, S, T and U**). We next examined pericytes from fibrotic kidneys of control and *Stat3* KO mice. For this, we isolated pericytes via FACS-sorting using NG2, CD140b positive and CD31 negative markers as shown in **Figure 5V**. Interestingly, we found that the gene expressions of the fibrotic factors including *Acta2*, *Col1a1*, *Fn1 Tgfb1*, and *Tgfbr1* was decreased in *Stat3* KO pericytes (**Figure 5V, W**). Taken together, these data indicate that STAT3 plays a direct role in regulating proliferation, cell detachment, cell spreading, cell migration and profibrotic signaling of pericytes leading to the development of fibrosis.

### STAT3 transcriptionally promotes *Collagen1a1* gene expression to execute fibrotic signaling *in vivo* and *in vitro*

To further probe how STAT3 regulates profibrotic signaling, we performed in silico analysis of promoter binding on *Collagen1a1* promoter. It was predicted that human and mouse *Collagen1a1* promoters contain consensus sequence for STAT3 binding (**Supplemental Figure 7**). Further, we used STAT3 ChIP-seq data from mouse kidney tissues (53). Re-analysis of ChIP sequencing data from WT kidneys shows that STAT3 binds on the *Collagen1a1* promoter but not on the *Fibronectin1* promoter (**Figure 6A, B**). To confirm the binding of STAT3 on the promoter of *Collagen1a1* gene, we performed ChIP assay in mouse kidneys and found that STAT3 binding was increased at day 14 post AA as compared to mice without AA treatment (**Figure 6C**). Moreover, we found a significant increase in the binding of STAT3 on the *Collagen1a1* promoter in our STAT3 gain of function pericytes cells as compared to control cells (**Figure 6D**). We then assessed the STAT3 transactivation in gain and loss of function pericytes cell models, using luciferase assay. We found a significant increase in luciferase activity in STAT3 gain of function models, while a significant decrease in loss of function models as compared to the respective control cells (**Figure 6E, F**). Thus, these data indicate that STAT3 directly regulates a key pro-fibrotic factor, *Collagen1a1*, by binding to its promoter *in vivo* and *in vitro*.

**Figure 6.**
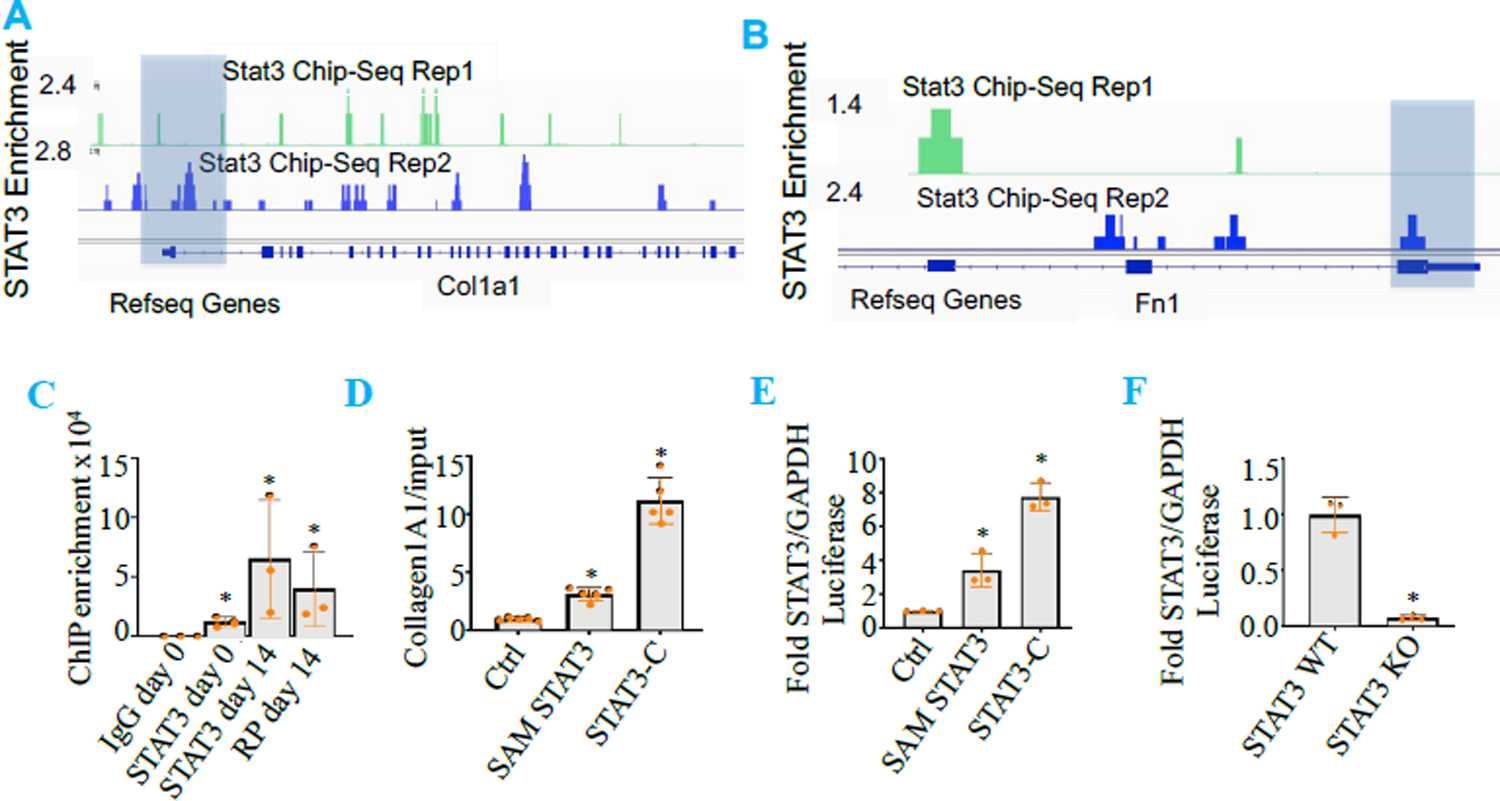
STAT3 binds on *Collagen1a1* promoter in mouse kidneys and in pericytes cell line to induce profibrotic signaling. (**A and B**) Analysis of previously published STAT3 ChIP sequencing data showing STAT3 binding on mouse *Collagen1a1* and Fibronectin 1 promoter region. (**C**) ChIP assay using anti-STAT3 antibody in normal and 14 days post AA injection in the mice kidneys to detect STAT3 binding on *Collagen1a1* promoter. Rabbit IgG and RNA polymerase (RP) antibodies are negative and positive controls. (**D**) ChIP assay using anti-STAT3 antibody in 10T1/2 cells with activated STAT3 to detect STAT3 binding on *Collagen1a1* promoter. Luciferase assay to detect STAT3 binding on *STAT3* consensus sequence in (**E**) STAT3 activated and (**F**) *Stat3* KO 10T1/2 cells. * represents p<0.05 as compared to control cells.

## DISCUSSION

Kidney fibrosis affects 10-15% of the population in the USA. While the functions of various cell types and their involvement has been documented but till now, the function of pericytes and its signaling pathway in tissue fibrosis remains poorly studied. Recent studies using mouse models of kidney, heart, and liver fibrosis by Kramann *et. al.* showed that sonic hedgehog mediated activation of Gli pathways controls fibrosis development (5, 54).

The origin of myofibroblasts in kidney fibrosis has been controversial. Whereas one group support the hypothesis that neither PDGFRβ positive nor NG2-positive cells contribute to the myofibroblast pool in the development of kidney fibrosis (17). Another group demonstrated that PDGFRβ positive cells do indeed contribute to the myofibroblast pool in kidney, lung, and heart fibrosis (55). Our data supports the hypothesis that PDGFRβ positive and NG2 positive pericytes detach from the endothelium and acquire myofibroblastic phenotype. The PDGFRβ positive myofibroblasts have been repeatedly described (16, 56, 57) but recent studies using *in vivo* genetic lineage tracing methods indicate that all myofibroblasts express PDGFRβ and that Gli1 positive cells do not acquire significant NG2 expression in fibrosis models (5).

Understanding the cellular and molecular signaling of the cell types involved in progression of kidney fibrosis is an important step towards the development of anti-fibrotic therapeutic strategies. STAT3 signaling has been implicated in organ fibrosis including kidney fibrosis (39, 58), where genetic ablation of *Stat3* from tubular epithelial cells protects mice from 5/6 nephrectomy induced kidney fibrosis by disrupting tubulointerstitial communication (39). In this previous report tubular cell function of STAT3 was well documented but the function of stromal cells remains unknown. Our data show that STAT3 in the stromal cells is crucial for the development of kidney fibrosis and genetic deletion of *Stat3* from Foxd1 positive cells protects mice from kidney fibrosis. Although delineating the quantitative contribution of specific cell sub-types from the stromal cells is needed to confirm the extent of protection, our results indicate 50% protection after *Stat3* deletion using the fibrotic assays such as IFTA scoring, semi-quantitative analyses of *Fibronectin1*, *Collagen1a1* and *Acta2* in two mechanistically different mouse models of kidney fibrosis. In addition to the mouse data, we found that STAT3 phosphorylation is significantly increased in interstitial cells of CKD patients. These findings are particularly important for the translational potential of STAT3 inhibition in patients with CKD.

Initiation of fibrosis is orchestrated by sequential activation of various cell types and one of the most common prominent pathways is inflammation. We and others have shown that DNA damage in AKI generates Reactive Oxygen Species (ROS), which triggers DNA damage signaling and activation of STAT3 pathways. In fact, STAT3 upregulation is associated with AKI (32, 59) and fibrosis development (38, 60–62). Persistent inflammation of tubules or interstitium initiates the progression of fibrosis. However, function of STAT3 in pericytes and its function/role in DNA damage and inflammation remains unclear. Our data for the first time show that *Stat3* depletion from stromal cells limits macrophage infiltration, restricts its inflammatory capabilities, and promotes M2 polarization, which helps to resolve inflammation. In accordance with these data, we found that *Stat3* depletion reduced DNA damage signaling and decreased proliferation of tubular epithelial cells and interstitial cells in the fibrotic kidneys of *Stat3* KO mice.

Myofibroblasts, a cell type that is well studied to lay matrix leading to fibrosis in chronic kidney disease, are differentiated from different cell types, pericytes being one of them (17, 57, 63). How pericytes differentiate into myofibroblasts is not well understood. We found that STAT3 is phosphorylated in tubular epithelial cells as well as in interstitial cells in human CKD kidneys and experimental mouse models of kidney fibrosis. Our data show that *Stat3* KO in stromal cells protects mice from FA or AA induced kidney fibrosis. Multiple models used in our study to increase STAT3 activation, including IL-6-mediated increase, CRISPR mediated transcriptional as well as mutation induced gain of function, all show that STAT3 is involved in proliferation, migration and increase in profibrotic signaling in pericyte-like cells. We found a significant increase in profibrotic signaling in STAT-C cells as compared to SAM STAT3 cells, which positively correlates with the amount of nuclear STAT3. Thus, confirming that STAT3 phosphorylation and its nuclear translocation directly regulate profibrotic signaling in pericyte-like cells. Stattic, a specific small molecular inhibitor of STAT3, inhibited IL-6 mediated proliferation, migration and profibrotic signaling and *Stat3* KO from the pericyte-like cells reduced the detachment, spreading proliferation and migration of these cells. Furthermore, pericytes isolated from *Stat3* KO fibrotic mice kidneys showed decreased expression of profibrotic signaling molecules, *Fibronectin1*, *Collagen1a1* and *Acta2*. Together, these data confirm a crucial function of STAT3 in the pericytes.

Binding of STAT3 on specific gene promoters decide the outcome of the cellular phenotype. For instance, STAT3 binding on *OCT4* and *NANOG* gene promoters regulates the stem cell proliferation (64), STAT3 binding on promoters of *Cyclin D1* and *c-MYC* cause cancer cell proliferation, controls cell cycle and deposits extracellular matrix (65), while binding of STAT3 on *Havcr1* promoter leads to tubular cell damage though activation of DNA damage induced inflammation (37, 60). Previously, we have shown that STAT3 binds on promoters of *Havcr1* and *fibrinogen α*, *β* as well as *γ* genes in AKI and fibrotic kidneys respectively (38, 60). Here, we found that STAT3 binds on the promoter of *Collagen1a1* gene in the mouse kidneys and pericyte-like cells and induces profibrotic signaling in the pericytes. The binding of STAT3 on specific DNA sequences during fibrosis progression needs further investigation to understand the dynamics of the whole genome occupancy. Thus, in this study we have identified a crucial role of STAT3 phosphorylation to increase profibrotic signaling in the stromal cells. These findings provide potential therapeutic interventions of pericytes using STAT3 as a therapeutic target (**Graphical Abstract**).

## STAR METHODS

### Key Resources table

#### Primary and secondary antibodies

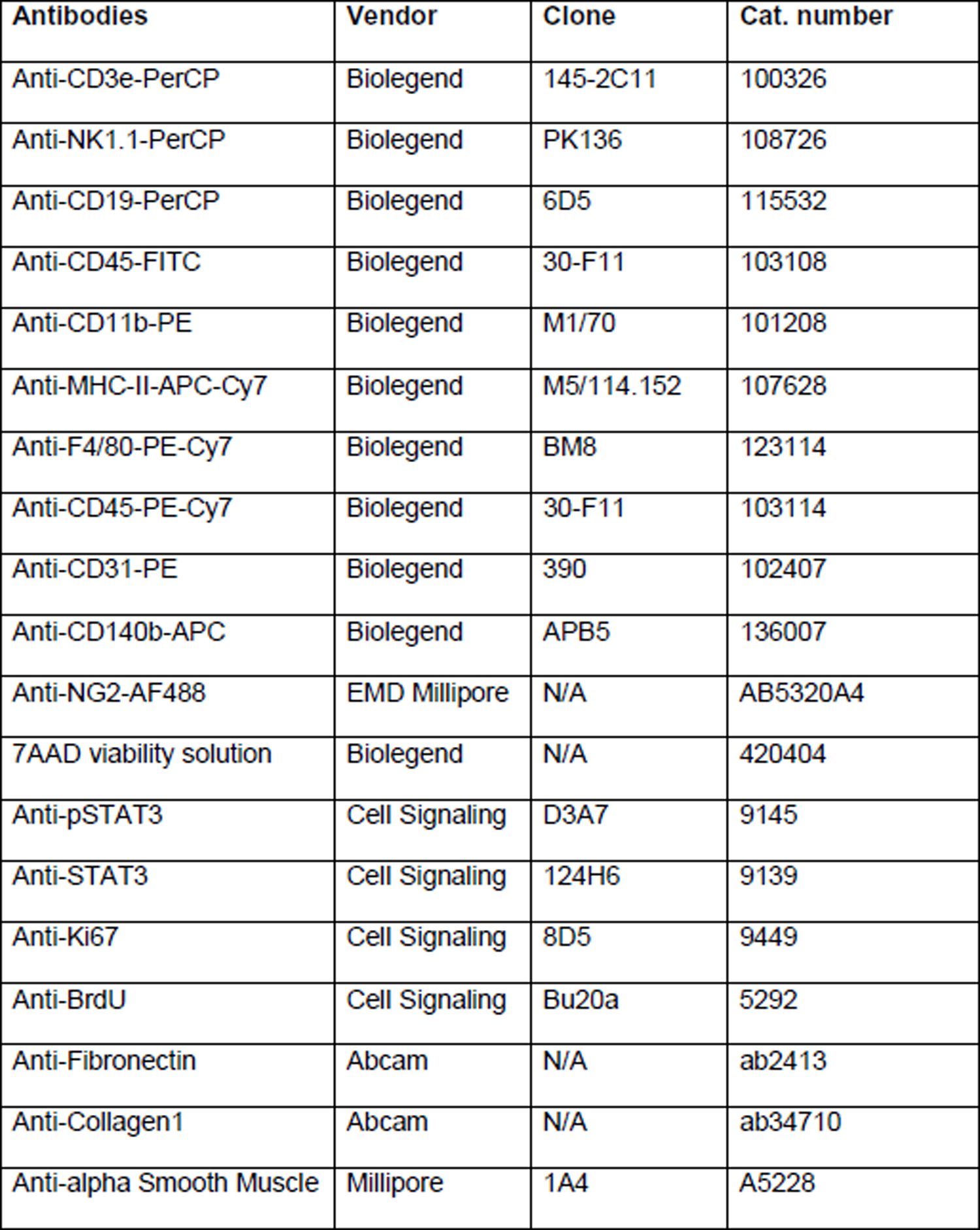
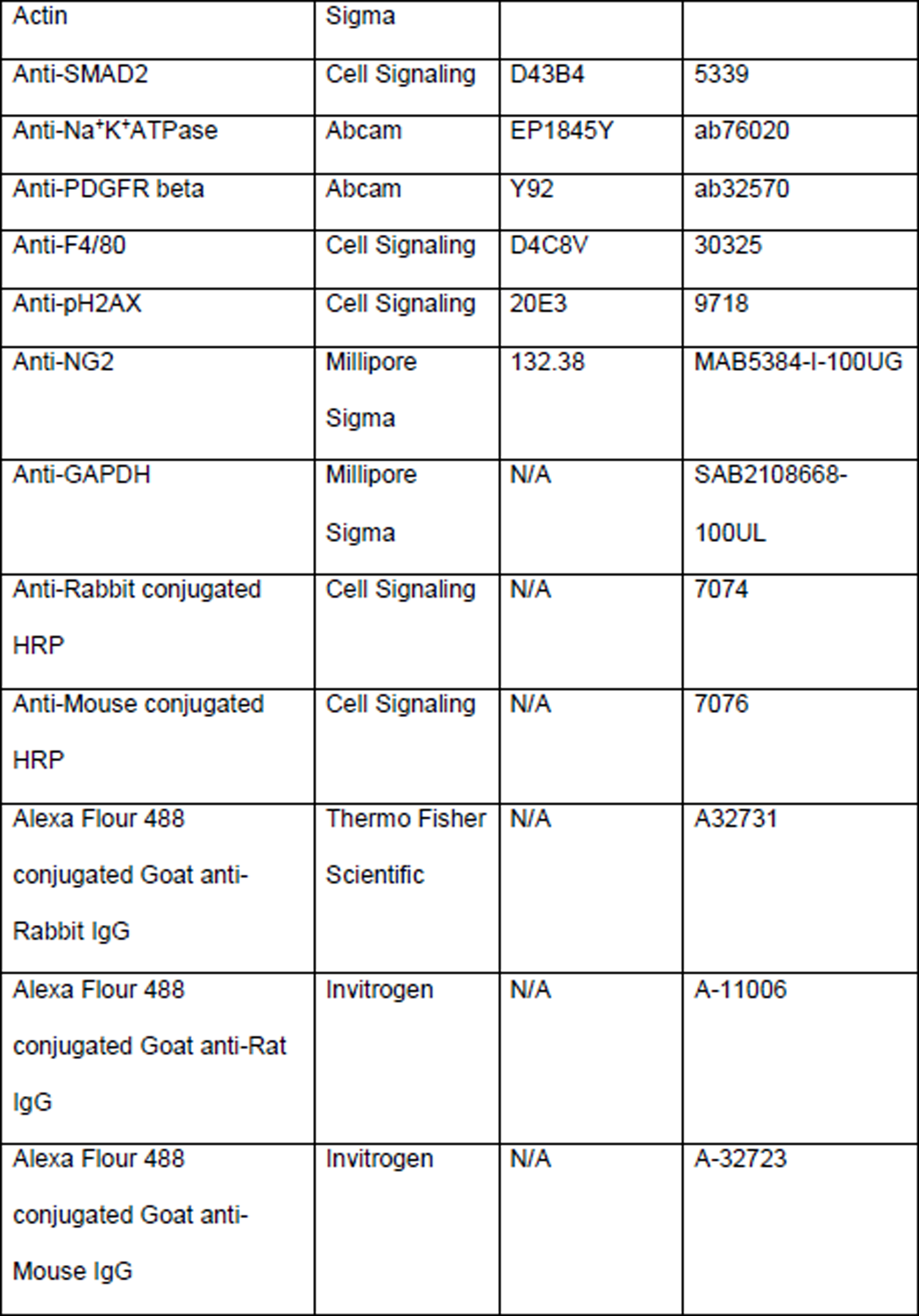
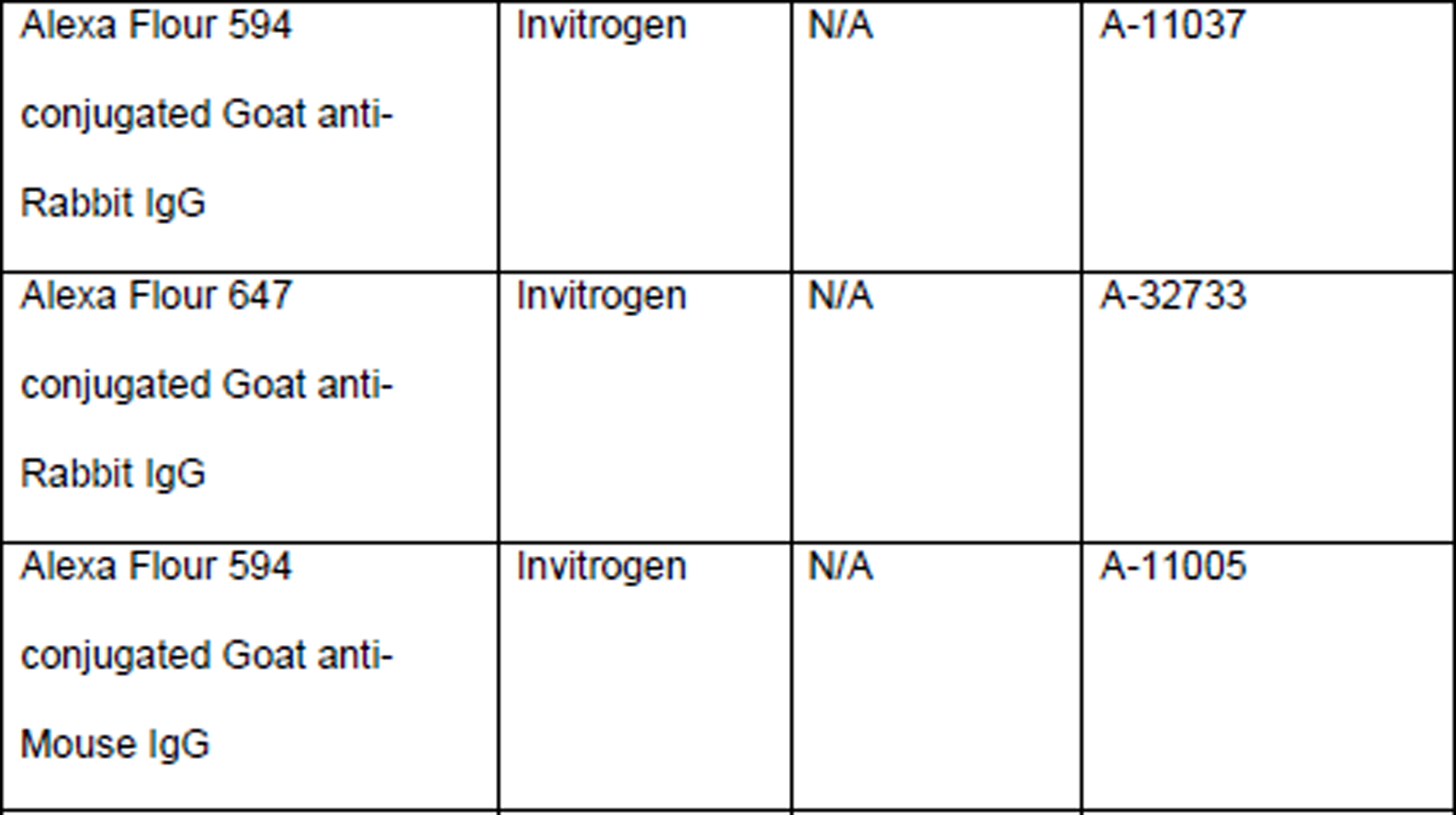

Primary and secondary antibodies:
| Antibodies | Vendor | Clone | Cat. number |
| --- | --- | --- | --- |
| Anti-CD3e-PerCP | Biolegend | 145-2C11 | 100326 |
| Anti-NK1.1-PerCP | Biolegend | PK136 | 108726 |
| Anti-CD19-PerCP | Biolegend | 6D5 | 115532 |
| Anti-CD45-FITC | Biolegend | 30-F11 | 103108 |
| Anti-CD11b-PE | Biolegend | M1/70 | 101208 |
| Anti-MHC-II-APC-Cy7 | Biolegend | M5/114.152 | 107628 |
| Anti-F4/80-PE-Cy7 | Biolegend | BM8 | 123114 |
| Anti-CD45-PE-Cy7 | Biolegend | 30-F11 | 103114 |
| Anti-CD31-PE | Biolegend | 390 | 102407 |
| Anti-CD140b-APC | Biolegend | APB5 | 136007 |
| Anti-NG2-AF488 | EMD Millipore | N/A | AB5320A4 |
| 7AAD viability solution | Biolegend | N/A | 420404 |
| Anti-pSTAT3 | Cell Signaling | D3A7 | 9145 |
| Anti-STAT3 | Cell Signaling | 124H6 | 9139 |
| Anti-Ki67 | Cell Signaling | 8D5 | 9449 |
| Anti-BrdU | Cell Signaling | Bu20a | 5292 |
| Anti-Fibronectin | Abcam | N/A | ab2413 |
| Anti-Collagen1 | Abcam | N/A | ab34710 |
| Anti-alpha Smooth Muscle | Millipore | 1A4 | A5228 |
| Actin | Sigma |  |  |
| Anti-SMAD2 | Cell Signaling | D43B4 | 5339 |
| Anti-Na<sup>+</sup>K<sup>+</sup>ATPase | Abcam | EP1845Y | ab76020 |
| Anti-PDGFR beta | Abcam | Y92 | ab32570 |
| Anti-F4/80 | Cell Signaling | D4C8V | 30325 |
| Anti-pH2AX | Cell Signaling | 20E3 | 9718 |
| Anti-NG2 | Millipore<br>Sigma | 132.38 | MAB5384-I-100UG |
| Anti-GAPDH | Millipore<br>Sigma | N/A | SAB2108668-100UL |
| Anti-Rabbit conjugated<br>HRP | Cell Signaling | N/A | 7074 |
| Anti-Mouse conjugated<br>HRP | Cell Signaling | N/A | 7076 |
| Alexa Flour 488<br>conjugated Goat anti-Rabbit IgG | Thermo Fisher<br>Scientific | N/A | A32731 |
| Alexa Flour 488<br>conjugated Goat anti-Rat<br>IgG | Invitrogen | N/A | A-11006 |
| Alexa Flour 488<br>conjugated Goat anti-Mouse IgG | Invitrogen | N/A | A-32723 |
| Alexa Flour 594<br>conjugated Goat anti-Rabbit IgG | Invitrogen | N/A | A-11037 |
| Alexa Flour 647<br>conjugated Goat anti-Rabbit IgG | Invitrogen | N/A | A-32733 |
| Alexa Flour 594<br>conjugated Goat anti-Mouse IgG | Invitrogen | N/A | A-11005 |

#### Chemicals and recombinant proteins

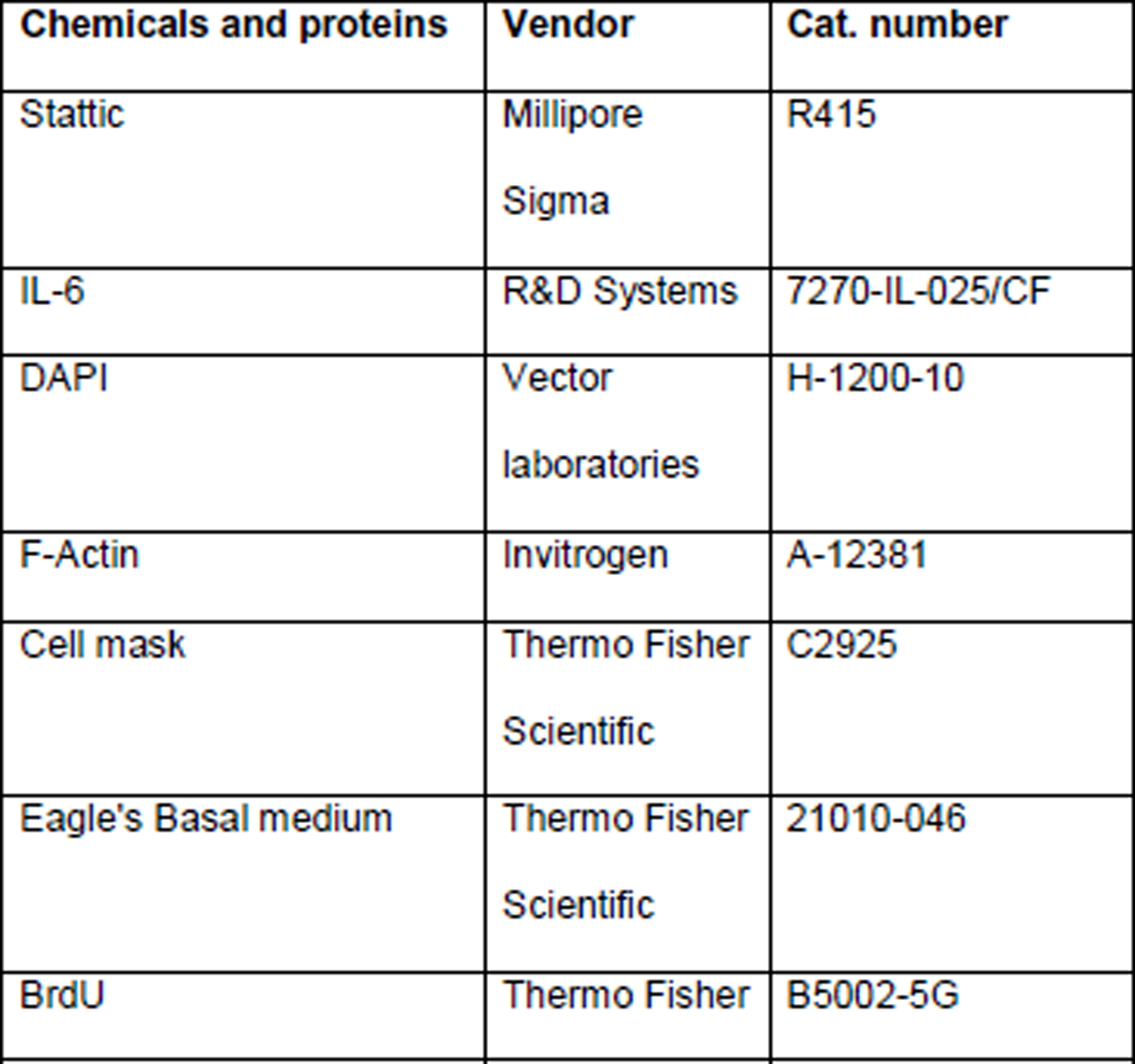
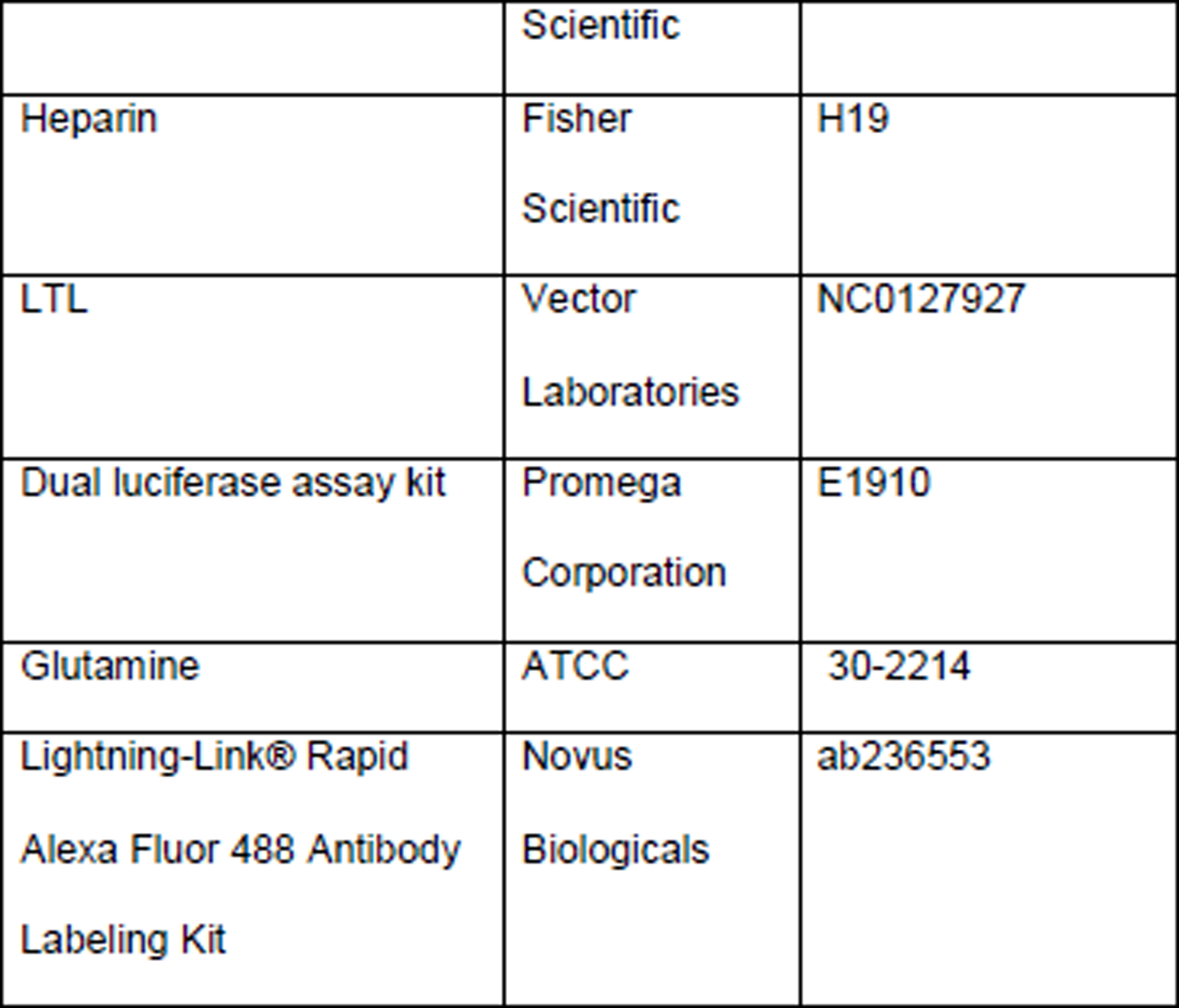

#### Mouse

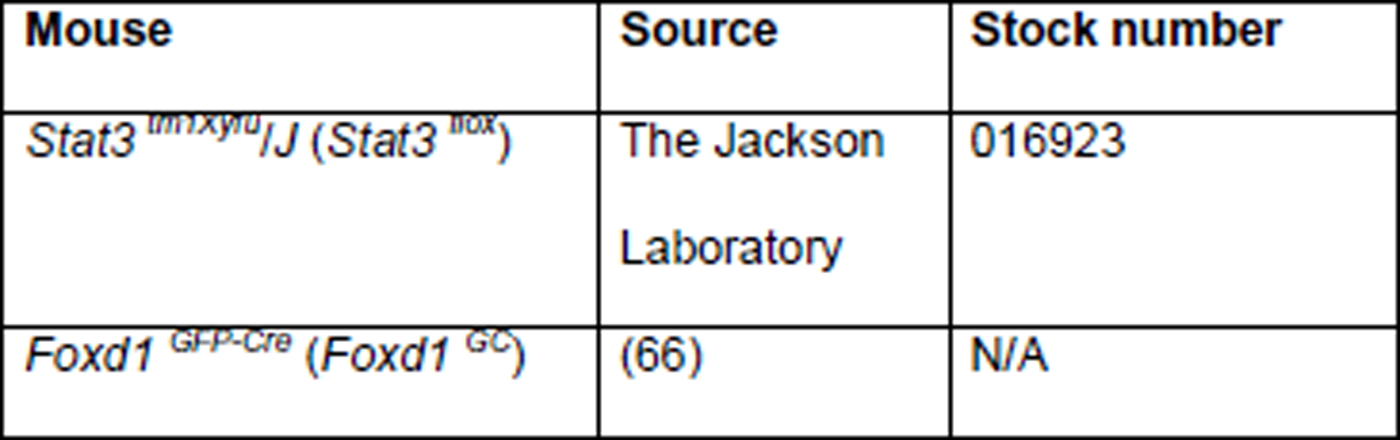

#### Software

Software:
| Name | Vendor | Identification<br>number |
| --- | --- | --- |
| GraphPad Prism 8 | GraphPad<br>Software | RRID:SCR_002798 |
| FIJI v1.0 | (67) | RRID:SCR_003070 |

## Human subjects

Human kidney biopsy sections were obtained through Brigham and Women’s Hospital’s pathology service core, and they were classified as patients without evidence of ATN (healthy, n=5), patients with evidence of Acute Tubular Necrosis (ATN) as AKI (n=5) and patients with Diabetic Nephropathy (DN) as CKD (n=5). The Institutional Review Board approved the protocol for extracting paraffin-embedded, formalin-fixed sections from patients with and without ATN and DN.

## Animals

Male C57BL/6J mice (age 8-10 weeks) were purchased from the Jackson Laboratory (Bar Harbor, ME). *Stat3 ^flox^ (Stat3 ^fl^*) mice were a kind gift from Dr. Kevin Haigis (Dana Farber Cancer Institute, Boston, MA) and *Foxd1 ^GC^* mice were a kind gift from Dr. Benjamin D. Humphreys (Washington University in St. Louis, MO). *Stat3* homozygous and *Foxd1* heterozygous mice were bred to generate *Stat3 ^fl/fl^ Foxd1 ^GC^* (*Stat3* KO) mice in *Foxd1* positive cell population. *Stat3* homozygous mice without *Foxd1 cre* allele named as *Stat3 ^fl/fl^* (control) mice were used as littermate controls. Mice were treated with single dose of either 250 mg/kg body weight of folic acid or 5 mg/kg body weight of aristolochic acid intraperitonially and blood was collected at acute and chronic phases of injuries. Mice were injected with 50 mg/kg body weight of BrdU, 3 hrs prior to sacrifice as previously described (68). Mice were sacrificed and kidneys were harvested at day 7 and 14 post treatments. All the experiments were conducted with the approval and guidelines of IACUC and Center for Comparative Medicine of Brigham and Women’s Hospital and were housed in a facility equipped with a 12-hour light/dark cycle.

## Estimation of BUN and sCr

BUN and sCr analysis were performed at O’Brien Centre AKI core at University of Alabama, AL and University of California, San Diego, CA, respectively.

## Fibrosis characterization

### Histology

Periodic Acid Schiff (PAS) stain and Mason’s Trichrome staining (MTS) was performed at Harvard rodent pathology core. Images were captured by a Nikon DS-Qi1Mc camera attached to a Nikon Eclipse 90i fluorescence microscope using an 40X or 60X objective using Nikon NIS elements AR version 3.2 software.

### Interstitial Fibrosis and tubular atrophy (IFTA) scoring

IFTA scores were performed by two pathologist and average of the scores were plotted.

## Chemical and antibodies

### Chemicals

IL-6 was purchased from R&D Systems (Minneapolis, MN) and resuspended as per manufacturer’s instructions. Stattic was purchased from Millipore Sigma (Burlington, MA) and dissolved in DMSO. Phalloidin (cat. no. R415) and Cell Tracker Green Dye (cat. no. C2925) was purchased from Thermo Fisher Scientific (Waltham, MA). Lotus Tetragonolobus Lectin (LTL) was purchased from Vector laboratories (cat. no. NC0127927).

### Antibodies

Anti-pSTAT3 Tyrosine 705 (cat. no. 9145) and anti-STAT3 (cat. no. 9139), anti-Ki67 (cat. no. 9449), anti-F4/80 (cat. no. 30325), pH2AX (cat. no. 9718), pSMAD2 (cat. no. 18338) were purchased from Cell Signaling Technology (Danvers, MA). Anti-Fibronectin (cat. no. ab2413), anti-Collagen1 (cat. no. ab34710) and anti-Na^+^K^+^ATPase (cat. no. ab76020) were purchased from Abcam (Cambridge, MA). Anti-α-SMA (cat. no. A5228) and anti-GAPDH (cat. no. SAB2108668-100UL) were purchased from Thermo Fisher Scientific. Horse Radish Peroxidase (HRP) conjugated anti-rabbit (cat. no. 7074) and anti-mouse (cat. no. 7076) antibodies were purchased from Cell Signaling Technology. Anti-Rabbit, anti-rat, or anti-mouse Alexa488, Alexa 594 and Alexa 647 conjugated antibodies were purchased from Thermo Fisher Scientific. For co-immunostaining using the same species antibodies, primary antibodies were conjugated using Lightning-Link® Rapid Alexa Fluor 488 Antibody Labeling Kit (Novus Biologicals, Centennial, CO 80112).

## Cell culture

10T1/2 a pericyte-like cells was purchased from ATCC (Manassas, VA) and maintained in BME medium with 10% FBS and 2 mg/ml sodium pyruvate. Cells were grown in a 37^0^C incubator with 5%CO_2_ and 37% relative humidity.

## CRISPR mediated gain and loss of function

### CRISPR mediated *Stat3* KO

sgRNAs were designed using Broad Institute/MIT CRISPR design tool (http://crispr.mit.edu) for the genomic location around transcription start site in the exon 1. sgRNAs were cloned into pSpCas9(BB)-2A-GFP plasmid (Addgene, Watertown, MA). Genomic DNA was extracted using the PureLink® Genomic DNA Mini Kit (cat. no. K182002, Thermo Fisher Scientific). PCR was performed using *Stat3* KO primers described in Table 1. The deletion was confirmed by DNA sequencing followed by western blotting of STAT3 (**Supplemental Figure 6A, B, C**).

Cells were transfected with the designed plasmid and incubated for 48 hrs with plasmids. Single cells positive for GFP were then sorted into a 96 well plate and were grown for 14 days with changing media every 4-5 days. Cells were then stained with pSTAT3 antibodies to confirm the expression levels of phosphorylated STAT3. Positive clones were selected for further studies.

**Table 1.**
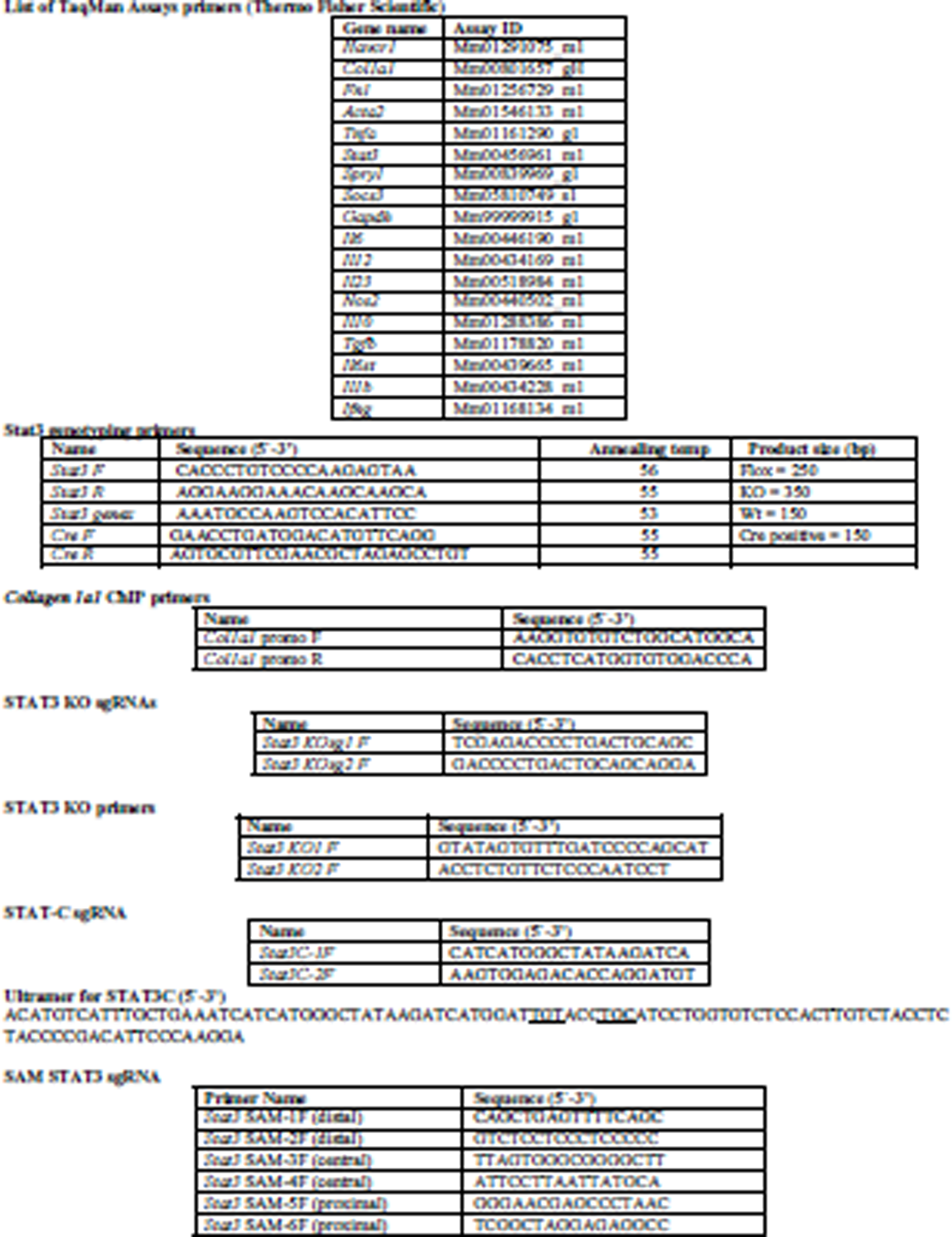

### CRISPR mediated Synergistic Activation Mediators (SAM)

For CRISPR mediated SAM, sgRNAs were designed for promoter region of *Stat3* (https://epd.epfl.ch//index.php) as discussed in the previous section and 4 residues were removed from 5’ end to make it a 16 nucleotides sgRNA. The 16-base pair sgRNA loses its capability to break the genomic DNA but retains the binding ability to specific DNA sequence (69). Two sgRNAs from proximal, central and distal promoter regions were selected and were cloned into the lenti-CRISPRV2 plasmid (Addgene). Lentivirus was produced in HEK293 cells by transfecting lenti-CRISPRV2, VSV-G, REV and GAG plasmids. The supernatant after incubation for 48 hrs was collected and centrifuged to get the clear supernatant containing viral particles.

Pericyte-like cells, 10T1/2 was transduced with the lentiviral particles. SAM activation plasmid containing MS2-p65-HSF1-VP64 was co-transfected in the cells along with the lentiviral particles. *Stat3* gene expression was confirmed by PCR amplification using TaqMan probes described in Table 1 and immunostaining with STAT3 Tyrosine 705 was performed at 72 hrs post transductions.

### CRISPR mediated mutation to generate STAT3-C

sgRNAs were designed by selecting the genomic location of Alanine662 and Asparagine664. sgRNAs were cloned into the pSpCas9(BB)-2A-GFP plasmid. Ultramers containing these two mutations to Cysteines were synthesized and co-transfected with the plasmid. GFP positive cells were sorted and were grown as single cell clones as described above. Immunostaining for pSTAT3 was performed to identify the cells with increased pSTAT3. List of sgRNAs and ultramers is shown in Table 1.

## Isolation of kidney pericytes and kidney resident macrophage using FACS

Control and *Stat3* KO mice were sacrificed at day 14 post AA treatment and perfused with ice cold PBS. Immediately, kidneys were harvested and digested with Iscove’s Modified Dulbecco’s Medium (IMDM) (Gibco) containing Collagenase D (2.5 mg/ml, Roche Diagnostics) and DNase I (1 mg/ml, Sigma-Aldrich) at 37 ^0^C for 45 min. Digested tissue was passed through a 70 µm cell strainer and centrifuged at 1600 rpm for 5 min at 4 ^0^C. Pellets were resuspend in 37% Percoll (GE Healthcare) and overlaid on top of 70% Percoll and centrifuged at 2000 rpm for 25 min at RT with no brake for gradient centrifugation. Mononuclear cells at the interphase were collected and washed twice with cold Hank’s Buffered Saline Solution (HBSS, Gibco). Samples were split into two, one for macrophage sorting and the other for pericyte sorting. Cells at the bottom of the tubes were also collected, washed twice with cold HBSS and included in the pericyte sorting. For pericyte sorting, cells were stained with 7-AAD, anti-CD45-PerCP, anti-CD3-PerCP, anti-NK1.1-PerCP, anti-CD19-PerCP, anti-CD31-PE, anti-CD140b-APC, anti-NG2-Alexa Fluor 488. For macrophage sort, cells were stained with 7-AAD, anti-CD45-PE-Cy7, anti-MHC II-APC-Cy7, anti-CD11b-PE, anti-F4/80-APC. Cells were then sorted using FACSAria IIu Cell Sorter (Becton Dickinson) using 100 µm nozzle. Collected cells were lysed with RLT buffer containing 1% β-mercaptoethanol.

## Scratch assay

10T1/2 cells were plated in a 6-well plate at a confluency of 80% and next day a scratch was created with a 200 μl microtip. Cells were washed three times with PBS and treated with IL-6 and stattic for 12 hrs. Cells were fixed and images were taken using Nikon Eclipse 90i microscope using 10X objective.

## Trans well migration assay

10T1/2 cells were plated 6-well plates and serum starved for 16 hrs. Ten thousand cells were then plated onto the upper well of 8 μm pore trans well inserts (BD Bioscience). The lower chamber contained either control medium, medium with IL-6 (100 ng/ml) or medium with IL-6 (100 ng/ml) and stattic (2.5 μM). Cells were incubated for additional 12 hrs. Using cotton swab the cells from the upper wells were removed and cells were fixed with 4% paraformaldehyde. Cells were then stained with cell mask green (1:10,000 dilution, Thermo Fisher Scientific) and were imaged after removing from the insert and fixing them on a glass slide using Nikon Eclipse 90i microscope using 10X objective.

## Western blotting

Western blotting was performed as previously described (36). Briefly, kidney tissues or cells were lysed using RIPA buffer containing protease and phosphatase inhibitors. Tissue was homogenized using homogenizer. Cell lysates were cleared using centrifugation and equal protein was loaded onto the gel. Protein was transferred onto the PVDF membrane and probed with specific antibodies. GAPDH or β-Actin were used as a loading control. Blots were developed using Luminata Forte HRP reagent using a Bio-Rad (Hercules, CA) or Syngene (Fredrick, MD) imaging systems. Quantification of blots was performed using Image lab for Bio-Rad or GeneSys for Syngene software and the data was normalized to the loading controls to calculate the fold change.

## RNA isolation, cDNA synthesis, and quantitative Real-Time PCR

Total RNA was isolated from cell pellets using the RNeasy Micro or Mini Kit (Qiagen, Germantown, MD) and first-strand cDNA synthesis was performed for each RNA sample from 0.5 to 1 μg of total RNA using TaqMan reverse transcription reagents.

### TaqMan RT-PCR

RNA from kidney tissues or cells was isolated using miRNeasy Mini Kit (Qiagen, Germantown, MD) as per manufacturer’s instruction. Five hundred nanograms of total RNA was converted into cDNA using High-Capacity cDNA Reverse Transcription Kits (Applied Biosystems, ThermoFisher Scientific) according to manufacturer’s protocol. Two microliters of five times diluted cDNA were used for PCR using TaqMan™ Universal PCR Master Mix with FAM labeled gene specific probes (ThermoFisher Scientific) in duplicate on a QuantStudio 7 thermal cycler (ThermoFisher Scientific). Fold change was calculated by delta Ct methods with *Gapdh* as reference gene and presented as fold change with respect to experimental controls.

### RT-PCR

For genotyping, DNA was extracted from mouse tail samples using Kappa Genotyping Kit (Roche, Basel, Switzerland) as per manufacturer’s instructions and RT-PCR was performed using specific primers described in Table 1. PCR products were resolved onto agarose gel and imaged using Bio-Rad gel imaging system ChemiDoc MP.

## Immunofluorescence staining

Immunofluorescence on OCT or paraffin tissue sections or cells were performed as previously described (36, 38, 68). Briefly, formalin fixed paraffin tissue sections were deparaffinized using xylene followed by antigen retrieval using citrate buffer. For the OCT block from PFA fixed tissue sections, PBS was added to the slides and antigen retrieval was performed using proteinase K treatment. For staining the cells, 4% PFA was used for fixation. Samples were permeabilized using 1% Triton X-100 for 15 min. Two percent normal sheep serum containing 5% BSA was used for blocking for 1 hr. Primary antibodies were incubated overnight at 4 ^0^C in five times diluted blocking buffer containing 0.1% Tween 20. After three washes of PBST (0.1% Tween 20), fluorescence labeled secondary antibodies (Thermo Fisher Scientific) were added and incubated for 1 h at room temperature. Slides were washed and mounted with medium containing DAPI (Vector laboratories). Images were captured on a Nikon Confocal Imaging system (Nikon C1 Eclipse, Nikon) using 60 X objective.

## BrdU incorporation assay

BrdU incorporation assay was performed as per manufacturer’s protocol (Abcam). Briefly, two thousand cells were plated in a 96 well plate and 2 hrs before endpoints, BrdU was added. Cells were fixed and anti-BrdU antibody was added and incubated at room temperature for 30 min. After washing, anti-mouse peroxidase IgG was added, and washing was repeated. TMB substrate was added and incubated in dark for 30 min at room temperature. Reaction was stopped by stop solution and plate was read at 450 nm. Control cells were referenced as 100% proliferation to plot the graphs.

## Cell detachment assay

Cells were plated on chambered slides and incubated for 1 h. Cells were then rotated on shaker for 15 min and washed thrice with PBS. Cells were fixed with 4% PFA and stained with phalloidin. Cells were then mounted with mounting medium containing DAPI. Images were captured using a Nikon confocal imaging system at 60x magnification. The number of cells present were counted, and graph were plotted.

## Cell spreading assay

Cells were plated on chambered slides and incubated for 3 h. Cells were fixed with 4% PFA and stained with phalloidin. Cells were then mounted with mounting medium containing DAPI. Images were captured using a Nikon confocal imaging system at 60x magnification. The area of cells was calculated using ImageJ software and graph were plotted with respect to control cells.

## In silico promoter analysis and ChIP sequencing analysis

### *In silico* STAT3 binding prediction

Promoter region of *Collagen1a1* and *Fibronectin1* gene were extracted from (https://epd.epfl.ch//index.php) and search was performed for STAT3 binding sites using (http://jaspar.genereg.net).

### ChIP sequencing analysis

ChIP data was analyzed from previously performed STAT3 ChIP sequencing in kidneys for promoter binding of *Collagen1a1* and *Fibronectin1* gene (GSM3176738, GSM3176739). Peaks on the log scale were shown as DNA binding sites for STAT3.

## Chromatin Immunoprecipitation (ChIP) assay

ChIP assay was performed as previously described (36). Briefly, 3-mm diameter of snap frozen kidney tissues or cells were treated with 1% formaldehyde/protease inhibitor DNA to crosslink proteins. Sonication was performed to get DNA fragments ranging from 200 to 1000 bp. Fifteen hundred μg protein was immunoprecipitated with 10 μg STAT3, normal rabbit IgG (negative control), or RNA polymerase (positive control) antibodies. Fifteen micrograms (1%) protein from each sample was frozen in 80°C as input. Protein G was used to pull down bound immune complex and DNA was isolated. Five-microliter samples were used for real time PCR using promoter-specific primers of *Collagen1a1* gene promoter (Table 1).

## Luciferase assay

Cells were plated in a 12 well plate and were co-transfected with Firefly *STAT3* luciferase and Renilla *Gapdh* luciferase (gift from Dr. Matthias Brock, Center of Experimental Rheumatology, University Hospital Zurich, Zurich, Switzerland) plasmids. After 24 hours cells were treated with IL-6 and stattic. Cells were harvested at 72 hrs post transfection and lysed using passive lysis buffer using Dual-Luciferase Reporter assay purchased from Promega (Madison, WI). Luciferase assay was performed as previously described (36). Ratio of Firefly and Renilla luciferase readings were plotted with respect to control cells.

## STATISTICAL ANALYSES

Statistical analyses were performed using an unpaired 2-tailed Student’s *t* test, or one- or two-way ANOVA as indicated. A *P* value of less than 0.05 was considered statistically significant. Data are generally presented as the mean ± SD. Please see individual figure captions for more details. GraphPad Prism 8.0 (GraphPad Software, Inc., La Jolla, CA) was used for all statistical analyses.

## ACKNOWLEDGEMENTS

This research was supported by a center grant to LH. AA is supported by a Career Development Grant from the American Heart Association (19CDA34780005). LZ is supported by the National Science Foundation, China. GM is supported by the grants from the NIH (R01 AI435801 and R01 AI151953). VS is supported by a R01 from the NCI.

## AUTHOR CONTRIBUTIONS

Conceptualization, AA, VS, AW, GM, DF, JB and LLH; Data curation, AA, LZ, SV, MF, KR, PC, AM, AB, MK, ST; Investigation, AA, LZ, SV; Visualization, AA, SV, MF, GM, ST; Methodology, AA, LZ, SV, GM, SJ, I-JC, KR, ST, VS; Writing, AA and LLH, Review and editing, all authors; Funding acquisition, AA, VS, JB, LLH.

## REFERENCES

1. Wynn TA. Common and unique mechanisms regulate fibrosis in various fibroproliferative diseases. J Clin Invest. 2007;117(3):524–9.

2. Zeisberg EM, Tarnavski O, Zeisberg M, Dorfman AL, McMullen JR, Gustafsson E, et al. Endothelial-to-mesenchymal transition contributes to cardiac fibrosis. Nat Med. 2007;13(8):952–61.

3. Quan TE, Cowper SE, and Bucala R. The role of circulating fibrocytes in fibrosis. Curr Rheumatol Rep. 2006;8(2):145–50.

4. Willis BC, duBois RM, and Borok Z. Epithelial origin of myofibroblasts during fibrosis in the lung. Proc Am Thorac Soc. 2006;3(4):377–82.

5. Kramann R, Schneider RK, DiRocco DP, Machado F, Fleig S, Bondzie PA, et al. Perivascular Gli1+ progenitors are key contributors to injury-induced organ fibrosis. Cell stem cell. 2015;16(1):51–66.

6. Mack M, and Yanagita M. Origin of myofibroblasts and cellular events triggering fibrosis. Kidney Int. 2015;87(2):297–307.

7. El Agha E, Kramann R, Schneider RK, Li X, Seeger W, Humphreys BD, et al. Mesenchymal Stem Cells in Fibrotic Disease. Cell stem cell. 2017;21(2):166–77.

8. E Oh, and Humphreys BD. Fibrotic Changes Mediating Acute Kidney Injury to Chronic Kidney Disease Transition. Nephron. 2017;137(4):264–7.

9. Meran S, and Steadman R. Fibroblasts and myofibroblasts in renal fibrosis. Int J Exp Pathol. 2011;92(3):158–67.

10. Grande MT, and Lopez-Novoa JM. Fibroblast activation and myofibroblast generation in obstructive nephropathy. Nat Rev Nephrol. 2009;5(6):319–28.

11. Schrimpf C, and Duffield JS. Mechanisms of fibrosis: the role of the pericyte. Curr Opin Nephrol Hypertens. 2011;20(3):297–305.

12. Lin SL, Chang FC, Schrimpf C, Chen YT, Wu CF, Wu VC, et al. Targeting endothelium-pericyte cross talk by inhibiting VEGF receptor signaling attenuates kidney microvascular rarefaction and fibrosis. Am J Pathol. 2011;178(2):911–23.

13. Humphreys BD, and DiRocco DP. Lineage-tracing methods and the kidney. Kidney Int. 2014;86(3):481–8.

14. Humphreys BD. Genetic tracing of the epithelial lineage during mammalian kidney repair. Kidney Int Suppl (2011). 2011;1(3):83-6.

15. Barnes JL, and Gorin Y. Myofibroblast differentiation during fibrosis: role of NAD(P)H oxidases. Kidney Int. 2011;79(9):944–56.

16. Humphreys BD, Lin SL, Kobayashi A, Hudson TE, Nowlin BT, Bonventre JV, et al. Fate tracing reveals the pericyte and not epithelial origin of myofibroblasts in kidney fibrosis. Am J Pathol. 2010;176(1):85–97.

17. LeBleu VS, Taduri G, O’Connell J, Teng Y, Cooke VG, Woda C, et al. Origin and function of myofibroblasts in kidney fibrosis. Nat Med. 2013;19(8):1047–53.

18. Yamashita T, Fujimiya M, Nagaishi K, Ataka K, Tanaka M, Yoshida H, et al. Fusion of bone marrow-derived cells with renal tubules contributes to renal dysfunction in diabetic nephropathy. FASEB journal: official publication of the Federation of American Societies for Experimental Biology. 2012;26(4):1559–68.

19. Tamaki K, Okuda S, Ando T, Iwamoto T, Nakayama M, and Fujishima M. TGF-beta 1 in glomerulosclerosis and interstitial fibrosis of adriamycin nephropathy. Kidney Int. 1994;45(2):525–36.

20. Yuan Q, Tan RJ, and Liu Y. Myofibroblast in Kidney Fibrosis: Origin, Activation, and Regulation. Adv Exp Med Biol. 2019;1165:253–83.

21. Costalonga EC, Fanelli C, Garnica MR, and Noronha IL. Adipose-Derived Mesenchymal Stem Cells Modulate Fibrosis and Inflammation in the Peritoneal Fibrosis Model Developed in Uremic Rats. Stem Cells Int. 2020;2020:3768718.

22. Wu F, Sun C, and Lu J. The Role of Chemokine Receptors in Renal Fibrosis. Rev Physiol Biochem Pharmacol. 2020.

23. Hewitson TD. Renal tubulointerstitial fibrosis: common but never simple. Am J Physiol Renal Physiol. 2009;296(6):F1239–44.

24. Strutz F, and Zeisberg M. Renal fibroblasts and myofibroblasts in chronic kidney disease. Journal of the American Society of Nephrology: JASN. 2006;17(11):2992–8.

25. Lin SL, Kisseleva T, Brenner DA, and Duffield JS. Pericytes and perivascular fibroblasts are the primary source of collagen-producing cells in obstructive fibrosis of the kidney. Am J Pathol. 2008;173(6):1617–27.

26. Duffield JS, and Humphreys BD. Origin of new cells in the adult kidney: results from genetic labeling techniques. Kidney Int. 2011;79(5):494–501.

27. Perry HM, Gorldt N, Sung SJ, Huang L, Rudnicka KP, Encarnacion IM, et al. Perivascular CD73(+) cells attenuate inflammation and interstitial fibrosis in the kidney microenvironment. Am J Physiol Renal Physiol. 2019;317(3):F658–F69.

28. Zhang P, Cai Y, Soofi A, and Dressler GR. Activation of Wnt11 by transforming growth factor-beta drives mesenchymal gene expression through non-canonical Wnt protein signaling in renal epithelial cells. J Biol Chem. 2012;287(25):21290–302.

29. Gewin L, and Zent R. How does TGF-beta mediate tubulointerstitial fibrosis? Semin Nephrol. 2012;32(3):228–35.

30. Toda N, Mukoyama M, Yanagita M, and Yokoi H. CTGF in kidney fibrosis and glomerulonephritis. Inflamm Regen. 2018;38:14.

31. Liu Z, Xue L, Liu Z, Huang J, Wen J, Hu J, et al. Tumor Necrosis Factor-Like Weak Inducer of Apoptosis Accelerates the Progression of Renal Fibrosis in Lupus Nephritis by Activating SMAD and p38 MAPK in TGF-beta1 Signaling Pathway. Mediators Inflamm. 2016;2016:8986451.

32. Nechemia-Arbely Y, Barkan D, Pizov G, Shriki A, Rose-John S, Galun E, et al. IL-6/IL-6R axis plays a critical role in acute kidney injury. Journal of the American Society of Nephrology: JASN. 2008;19(6):1106–15.

33. Donate-Correa J, Martin-Nunez E, Muros-de-Fuentes M, Mora-Fernandez C, and Navarro-Gonzalez JF. Inflammatory cytokines in diabetic nephropathy. J Diabetes Res. 2015;2015:948417.

34. Okamoto M, Lee C, and Oyasu R. Interleukin-6 as a paracrine and autocrine growth factor in human prostatic carcinoma cells in vitro. Cancer Res. 1997;57(1):141–6.

35. Leu CM, Wong FH, Chang C, Huang SF, and Hu CP. Interleukin-6 acts as an antiapoptotic factor in human esophageal carcinoma cells through the activation of both STAT3 and mitogen-activated protein kinase pathways. Oncogene. 2003;22(49):7809–18.

36. Ajay AK, Kim TM, Ramirez-Gonzalez V, Park PJ, Frank DA, and Vaidya VS. A bioinformatics approach identifies signal transducer and activator of transcription-3 and checkpoint kinase 1 as upstream regulators of kidney injury molecule-1 after kidney injury. J Am Soc Nephrol. 2014;25(1):105–18.

37. Collier JB, and Schnellmann RG. Extracellular Signal-Regulated Kinase 1/2 Regulates Mouse Kidney Injury Molecule-1 Expression Physiologically and Following Ischemic and Septic Renal Injury. J Pharmacol Exp Ther. 2017;363(3):419–27.

38. Craciun FL, Ajay AK, Hoffmann D, Saikumar J, Fabian SL, Bijol V, et al. Pharmacological and genetic depletion of fibrinogen protects from kidney fibrosis. Am J Physiol Renal Physiol. 2014;307(4):F471–84.

39. Bienaime F, Muorah M, Yammine L, Burtin M, Nguyen C, Baron W, et al. Stat3 Controls Tubulointerstitial Communication during CKD. Journal of the American Society of Nephrology: JASN. 2016;27(12):3690–705.

40. Nagalakshmi VK, Li M, Shah S, Gigliotti JC, Klibanov AL, Epstein FH, et al. Changes in cell fate determine the regenerative and functional capacity of the developing kidney before and after release of obstruction. Clin Sci (Lond). 2018;132(23):2519–45.

41. Li P, Zhou Y, Goodwin AJ, Cook JA, Halushka PV, Zhang XK, et al. Fli-1 Governs Pericyte Dysfunction in a Murine Model of Sepsis. J Infect Dis. 2018;218(12):1995–2005.

42. Saito Y, Yamanaka S, Fujimoto T, Tajiri S, Matsumoto N, Takamura T, et al. Mesangial cell regeneration from exogenous stromal progenitor by utilizing embryonic kidney. Biochem Biophys Res Commun. 2019;520(3):627–33.

43. Kida Y, and Duffield JS. Pivotal role of pericytes in kidney fibrosis. Clin Exp Pharmacol Physiol. 2011;38(7):467–73.

44. Kramann R, and Humphreys BD. Kidney pericytes: roles in regeneration and fibrosis. Semin Nephrol. 2014;34(4):374–83.

45. Wang N, Deng Y, Liu A, Shen N, Wang W, Du X, et al. Novel Mechanism of the Pericyte-Myofibroblast Transition in Renal Interstitial Fibrosis: Core Fucosylation Regulation. Sci Rep. 2017;7(1):16914.

46. Wu CF, Chiang WC, Lai CF, Chang FC, Chen YT, Chou YH, et al. Transforming growth factor beta-1 stimulates profibrotic epithelial signaling to activate pericyte-myofibroblast transition in obstructive kidney fibrosis. Am J Pathol. 2013;182(1):118–31.

47. Artaza JN, Singh R, Ferrini MG, Braga M, Tsao J, and Gonzalez-Cadavid NF. Myostatin promotes a fibrotic phenotypic switch in multipotent C3H 10T1/2 cells without affecting their differentiation into myofibroblasts. J Endocrinol. 2008;196(2):235–49.

48. Fabian SL, Penchev RR, St-Jacques B, Rao AN, Sipila P, West KA, et al. Hedgehog-Gli pathway activation during kidney fibrosis. Am J Pathol. 2012;180(4):1441–53.

49. DiRocco DP, Kobayashi A, Taketo MM, McMahon AP, and Humphreys BD. Wnt4/beta-catenin signaling in medullary kidney myofibroblasts. Journal of the American Society of Nephrology: JASN. 2013;24(9):1399–412.

50. Ding J, Su S, You T, Xia T, Lin X, Chen Z, et al. Serum interleukin-6 level is correlated with the disease activity of systemic lupus erythematosus: a meta-analysis. Clinics (Sao Paulo). 2020;75:e1801.

51. Chen TK, Estrella MM, Appel LJ, Coresh J, Luo S, Obeid W, et al. Serum levels of interleukin (IL)-6, IL-8 and IL-10 and risks of end-stage kidney disease and mortality. Nephrol Dial Transplant. 2020.

52. Magno AL, Herat LY, Carnagarin R, Schlaich MP, and Matthews VB. Current Knowledge of IL-6 Cytokine Family Members in Acute and Chronic Kidney Disease. Biomedicines. 2019;7(1).

53. Lee HK, Willi M, Shin HY, Liu C, and Hennighausen L. Progressing super-enhancer landscape during mammary differentiation controls tissue-specific gene regulation. Nucleic Acids Res. 2018;46(20):10796–809.

54. Kramann R, Fleig SV, Schneider RK, Fabian SL, DiRocco DP, Maarouf O, et al. Pharmacological GLI2 inhibition prevents myofibroblast cell-cycle progression and reduces kidney fibrosis. J Clin Invest. 2015;125(8):2935–51.

55. Henderson NC, Arnold TD, Katamura Y, Giacomini MM, Rodriguez JD, McCarty JH, et al. Targeting of alphav integrin identifies a core molecular pathway that regulates fibrosis in several organs. Nat Med. 2013;19(12):1617–24.

56. Asada N, Takase M, Nakamura J, Oguchi A, Asada M, Suzuki N, et al. Dysfunction of fibroblasts of extrarenal origin underlies renal fibrosis and renal anemia in mice. J Clin Invest. 2011;121(10):3981–90.

57. Kramann R, DiRocco DP, and Humphreys BD. Understanding the origin, activation and regulation of matrix-producing myofibroblasts for treatment of fibrotic disease. J Pathol. 2013;231(3):273–89.

58. Kasembeli MM, Bharadwaj U, Robinson P, and Tweardy DJ. Contribution of STAT3 to Inflammatory and Fibrotic Diseases and Prospects for its Targeting for Treatment. Int J Mol Sci. 2018;19(8).

59. Dube S, Matam T, Yen J, Mang HE, Dagher PC, Hato T, et al. Endothelial STAT3 Modulates Protective Mechanisms in a Mouse Ischemia-Reperfusion Model of Acute Kidney Injury. J Immunol Res. 2017;2017:4609502.

60. Ajay AK, Kim TM, Ramirez-Gonzalez V, Park PJ, Frank DA, and Vaidya VS. A bioinformatics approach identifies signal transducer and activator of transcription-3 and checkpoint kinase 1 as upstream regulators of kidney injury molecule-1 after kidney injury. Journal of the American Society of Nephrology: JASN. 2014;25(1):105–18.

61. Pace J, Paladugu P, Das B, He JC, and Mallipattu SK. Targeting STAT3 signaling in kidney disease. Am J Physiol Renal Physiol. 2019;316(6):F1151–F61.

62. Pang M, Ma L, Gong R, Tolbert E, Mao H, Ponnusamy M, et al. A novel STAT3 inhibitor, S3I-201, attenuates renal interstitial fibroblast activation and interstitial fibrosis in obstructive nephropathy. Kidney Int. 2010;78(3):257-68.

63. Campanholle G, Ligresti G, Gharib SA, and Duffield JS. Cellular mechanisms of tissue fibrosis. 3. Novel mechanisms of kidney fibrosis. Am J Physiol Cell Physiol. 2013;304(7):C591-603.

64. Do DV, Ueda J, Messerschmidt DM, Lorthongpanich C, Zhou Y, Feng B, et al. A genetic and developmental pathway from STAT3 to the OCT4-NANOG circuit is essential for maintenance of ICM lineages in vivo. Genes Dev. 2013;27(12):1378–90.

65. Zhang F, Li C, Halfter H, and Liu J. Delineating an oncostatin M-activated STAT3 signaling pathway that coordinates the expression of genes involved in cell cycle regulation and extracellular matrix deposition of MCF-7 cells. Oncogene. 2003;22(6):894–905.

66. Kobayashi A, Mugford JW, Krautzberger AM, Naiman N, Liao J, and McMahon AP. Identification of a multipotent self-renewing stromal progenitor population during mammalian kidney organogenesis. Stem Cell Reports. 2014;3(4):650–62.

67. Schindelin J, Arganda-Carreras I, Frise E, Kaynig V, Longair M, Pietzsch T, et al. Fiji: an open-source platform for biological-image analysis. Nat Methods. 2012;9(7):676–82.

68. Ajay AK, Saikumar J, Bijol V, and Vaidya VS. Heterozygosity for fibrinogen results in efficient resolution of kidney ischemia reperfusion injury. PLoS One. 2012;7(9):e45628.

69. Dahlman JE, Abudayyeh OO, Joung J, Gootenberg JS, Zhang F, and Konermann S. Orthogonal gene knockout and activation with a catalytically active Cas9 nuclease. Nat Biotechnol. 2015;33(11):1159–61.

