## Supplementary figures and images for "Stromal-cell deletion of STAT3 protects mice from kidney fibrosis by inhibiting pericytes trans-differentiation and migration"

### Supplemental Figure 1

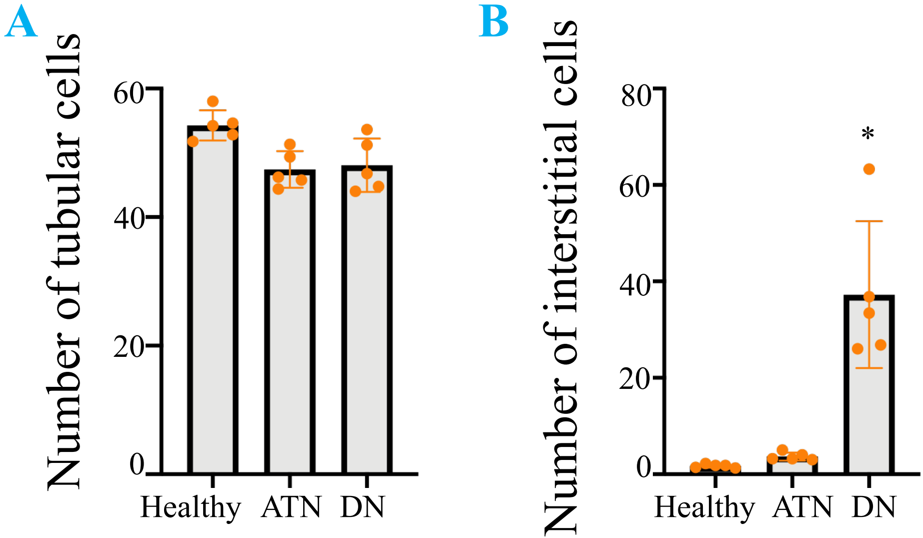


Supplementary Figure 1

### Supplemental Figure 2

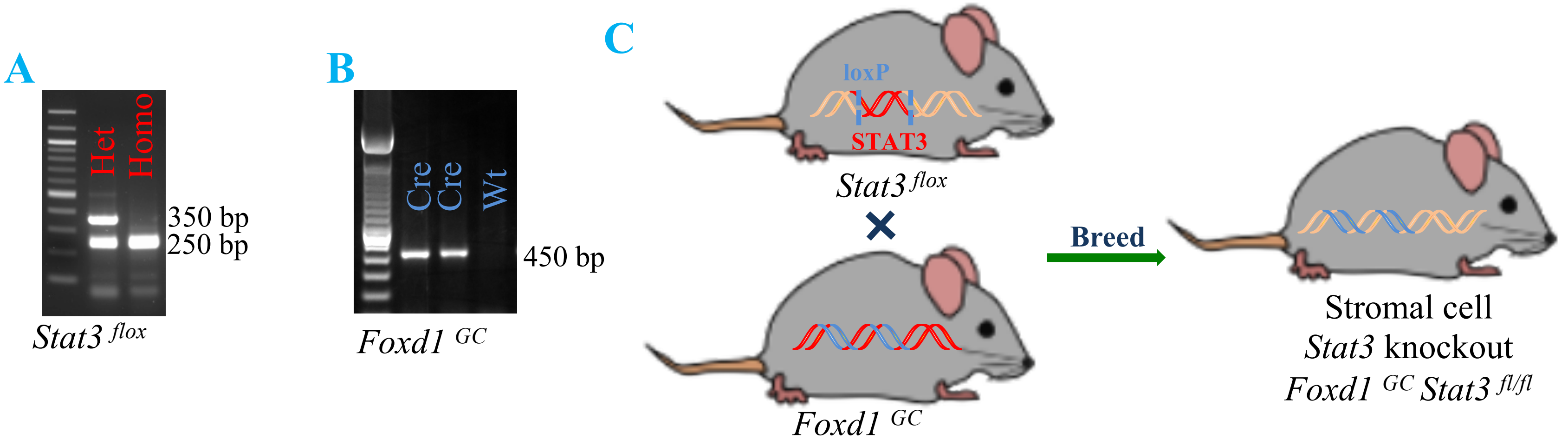


Supplementary Figure 2

### Supplemental Figure 3

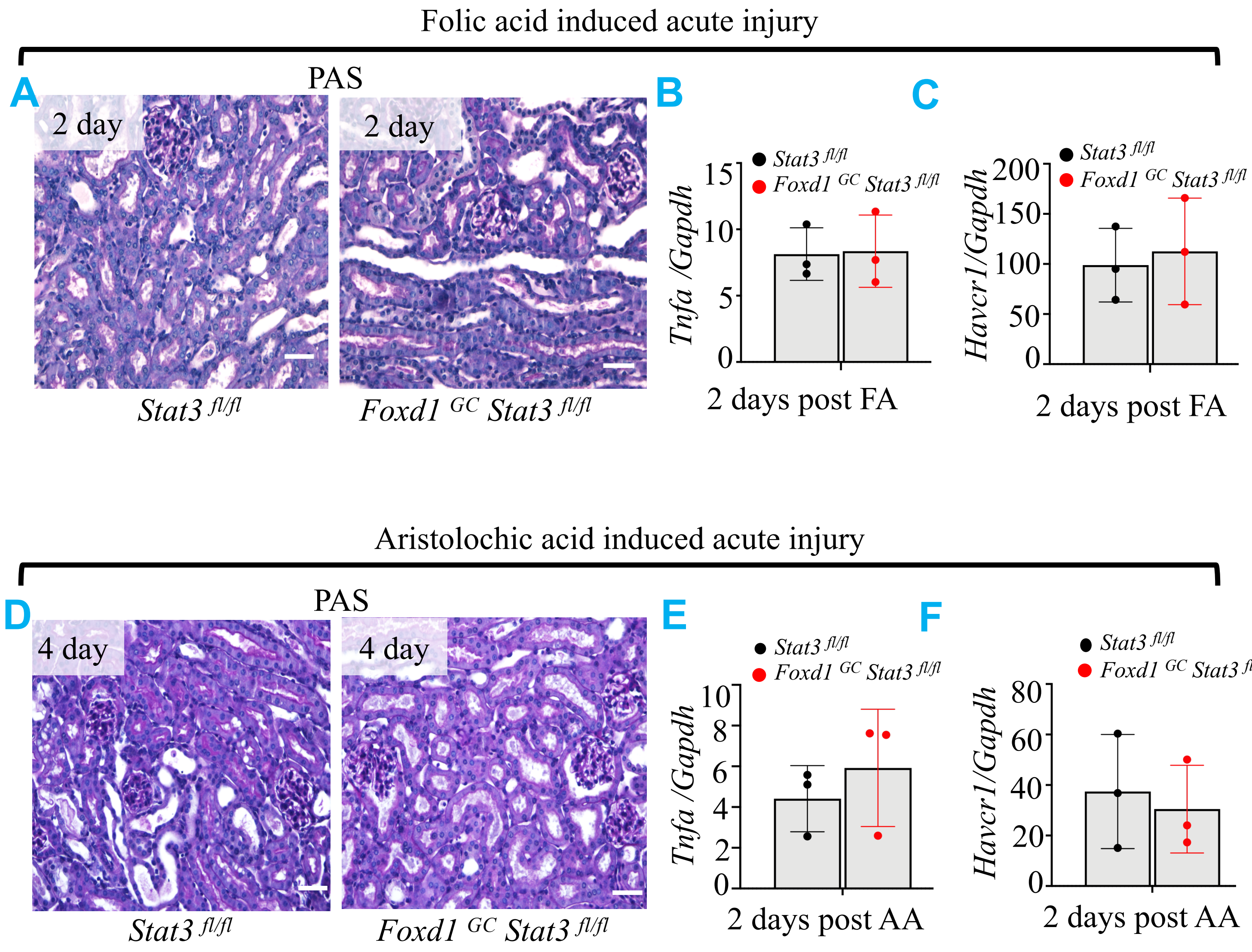


Supplementary Figure 3

### Supplemental Figure 4

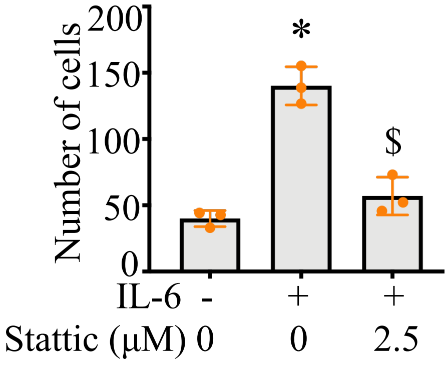


Supplementary Figure 4

### Supplemental Figure 5

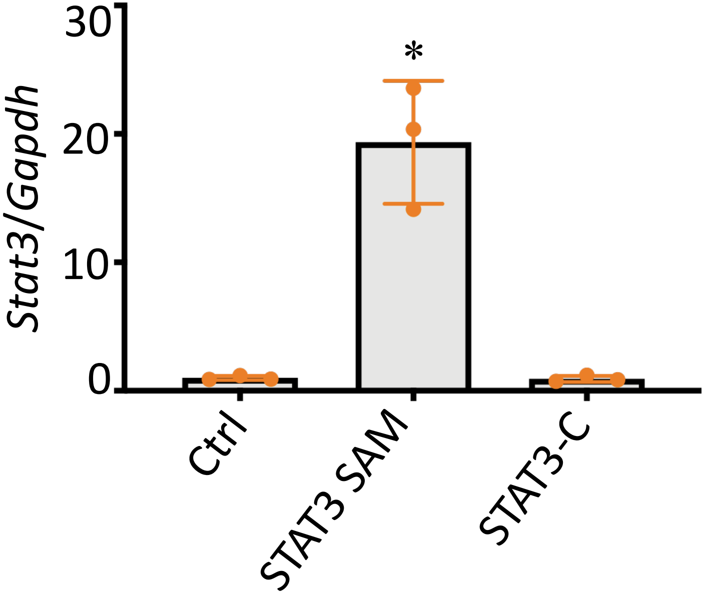


Supplementary Figure 5

### Supplemental Figure 6

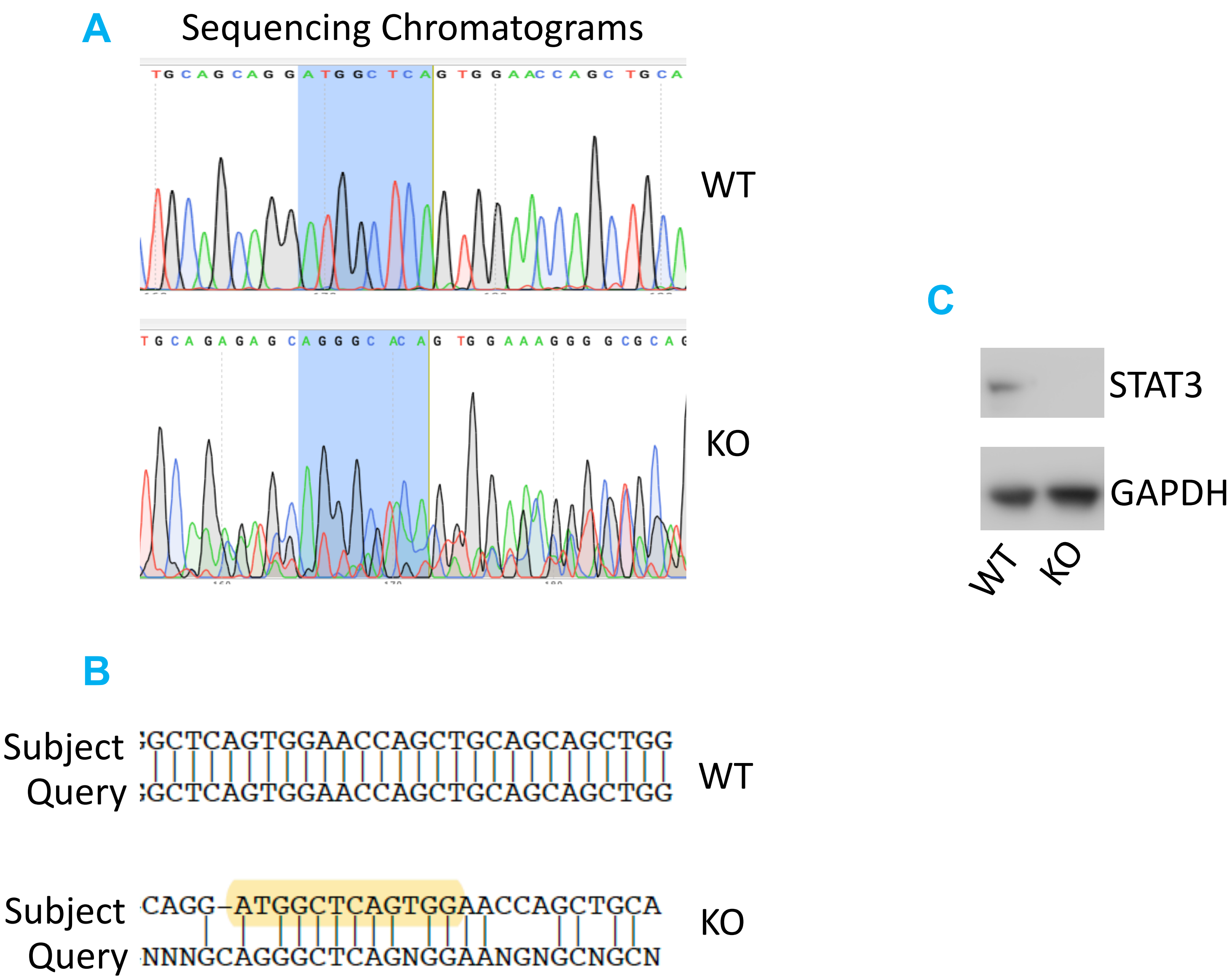


Supplementary Figure 6
