## Supplemental Figure 7 for "Stromal-cell deletion of STAT3 protects mice from kidney fibrosis by inhibiting pericytes trans-differentiation and migration"

STAT3 binding sites on human COL1A1 promoter

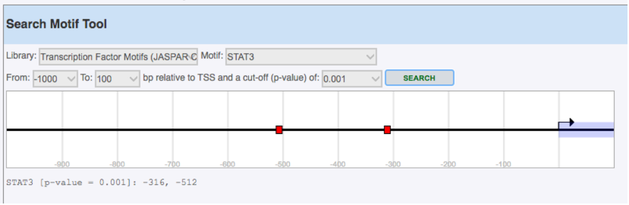


STAT3 binding sites on mouse Col1a1 promoter

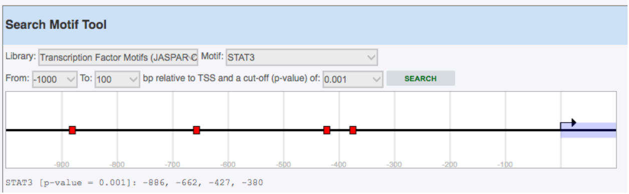


Supplementary Figure 7
