## Supplementary data for "Stromal-cell deletion of STAT3 protects mice from kidney fibrosis by inhibiting pericytes trans-differentiation and migration"

**Supplemental data**

**Methods:**

1. **DNA sequence analysis:** DNA sequencing data was viewed using SnapGene Viewer to obtain the chromatograms. Sequences were aligned using pairwise alignment tool using Needle ([https://www.ebi.ac.uk/Tools/psa/emboss_needle](https://www.ebi.ac.uk/Tools/psa/emboss_needle/)).
2. **PAS staining:** PAS staining was performed at Rodent Histology Core, Harvard Medical School, Boston, MA.

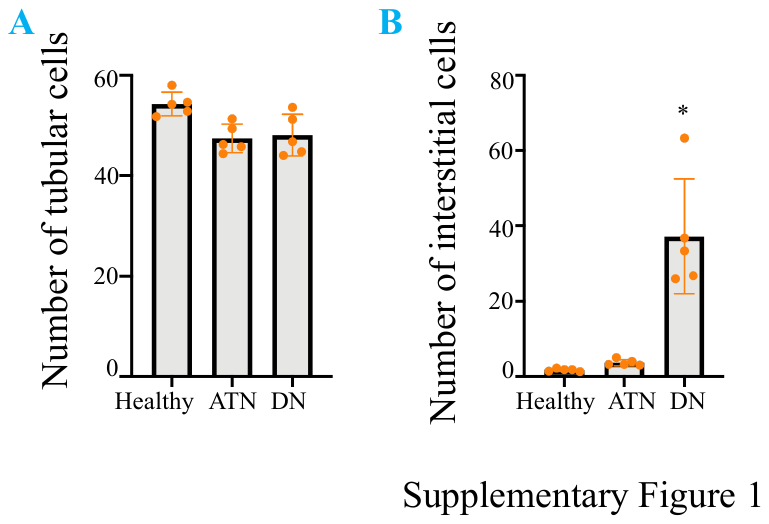

**Supplemental Figure 1:** Total number of DAPI positive nuclei from (A) tubular cells and (B) interstitial cells in healthy, ATN and DN human kidney tissues. * Represents p ≤ 0.05.

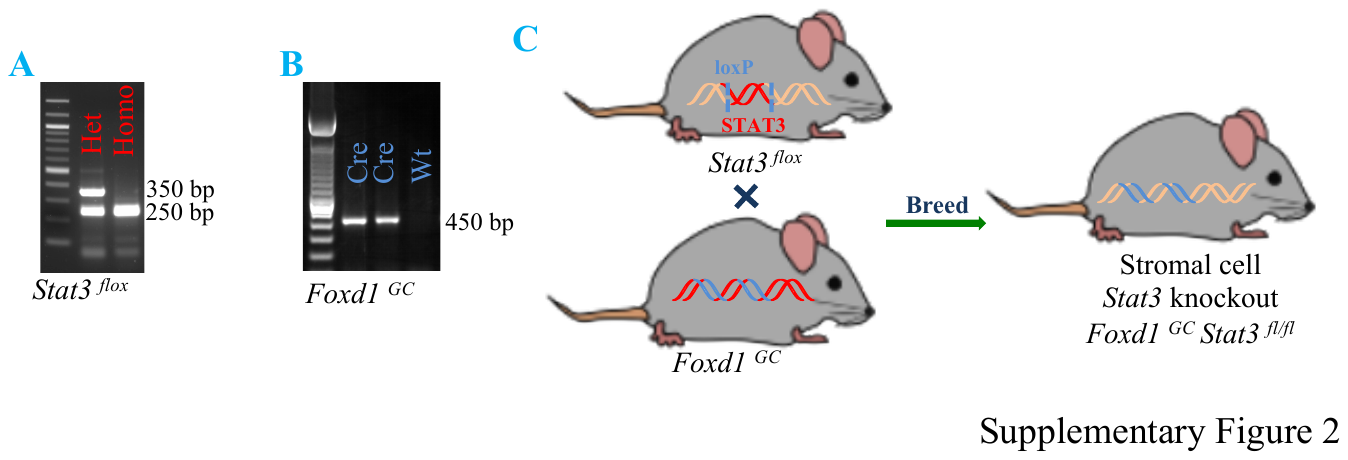
**Supplemental Figure 2:** Development of *Stat3* KO in *Foxd1* population. (A) Genotyping for *Stat3 flox* allele. (B) Genotyping for *Cre* recombinase. (C) Breeding strategy for *Stat3* *flox* and *Foxd1 Cre* to first generated heterozygous *Stat3* and *Foxd1 Cre* F1 generation (not shown). Backcrossing these F1 with homozygous *Stat3 flox* to generate homozygous *Stat3 flox* and heterozygous *Foxd1 Cre*.

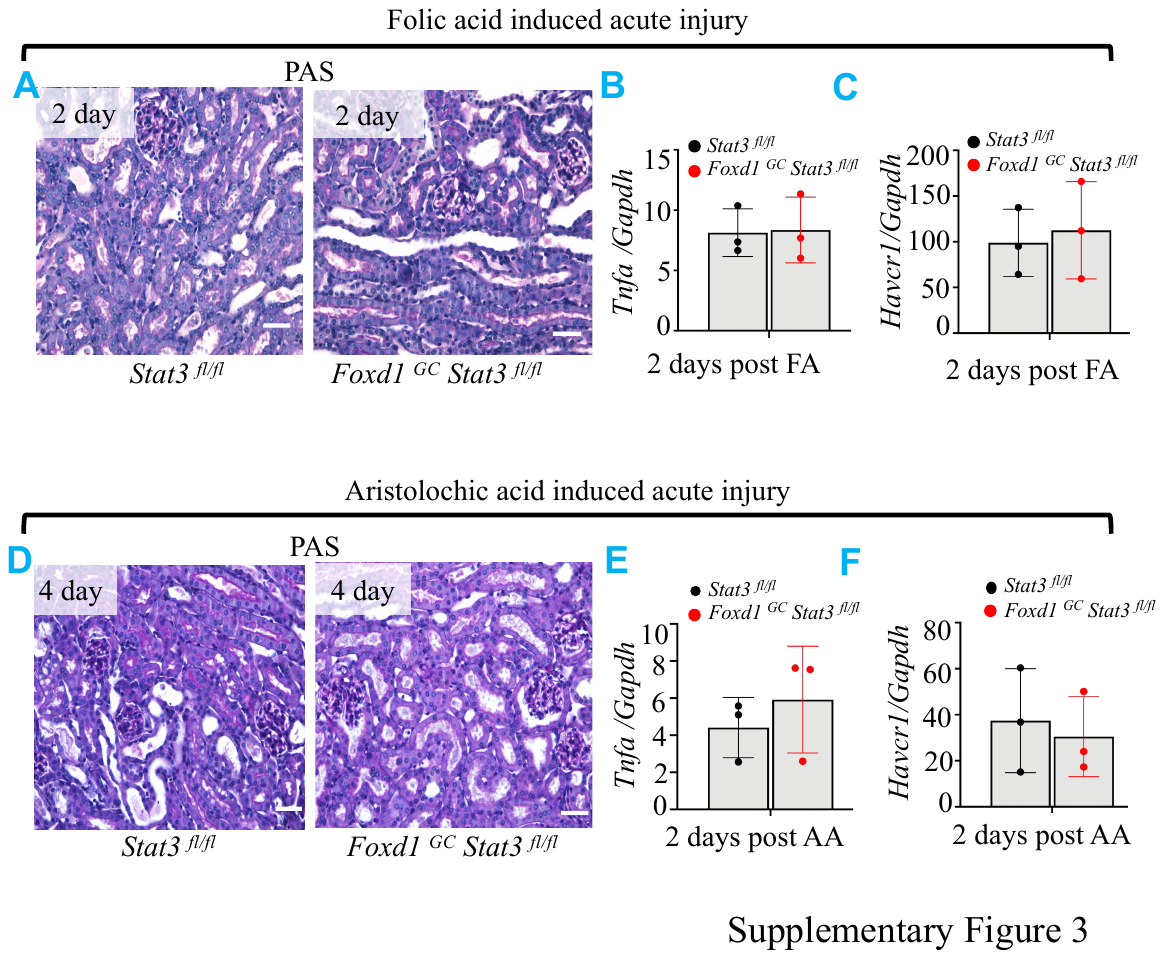

**Supplemental Figure 3:** Analysis of AKI phase of FA or AA-induced injury in *Stat3* control and *Stat3* KO mice kidneys. (A, D) Histological analysis using PAS staining. (B, C, E, F) Gene expression analysis of *Tnfa* and *Havcr1* by qRT-PCR.

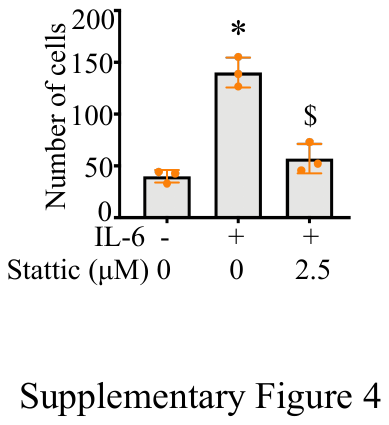

**Supplemental Figure 4:** Total number of cells were counted by counting DAPI positive nuclei following IL-6 and/or stattic treatments. * Represents p ≤ 0.05 as compared to control cells and $ represents p ≤ 0.05 as compared to IL-6 treated cells.

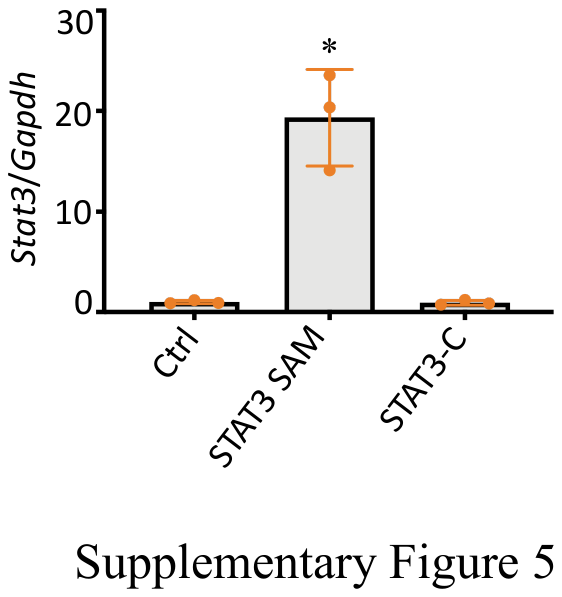

**Supplemental Figure 5:** Quantitative RT-PCR to detect *Stat3* mRNA in SAM STAT3 and STAT3-C cells. * Represents p ≤ 0.05 as compared to control cells.

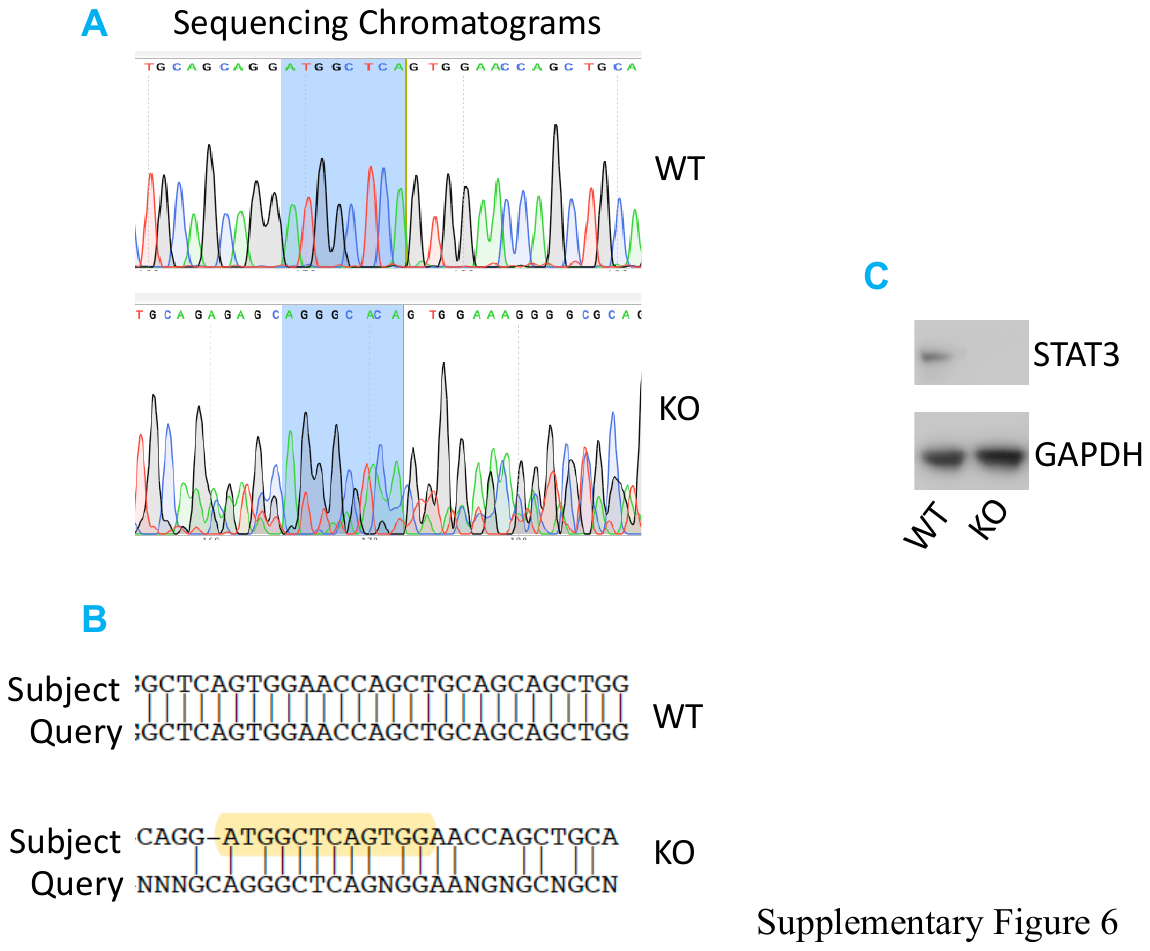

**Supplemental Figure 6:** Confirmation of *Stat3* KO in 10T1/2 cells. (A) Chromatogram of single cell clone and wildtype cells following DNA sequencing. (B) Sequence alignment of DNA sequencing of single cell clones with wildtype cells. (C) STAT3 immunoblotting in *Stat3* WT and KO cells. GAPDH served as a reference protein.

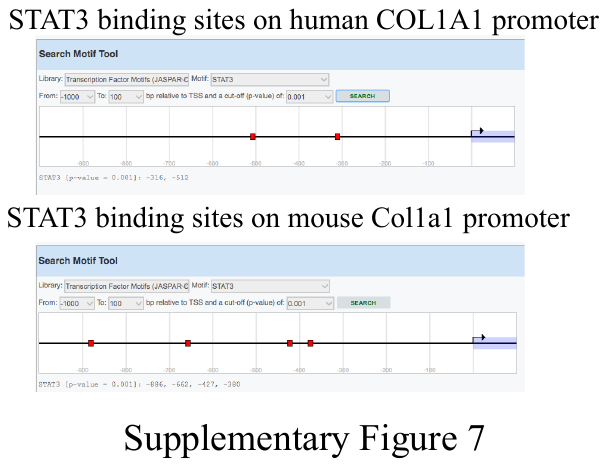

**Supplemental Figure 7:** (**A**) *In silico* promoter analysis for STAT3 binding sites on human and mouse *Collagen1a1* promoter. STAT3 binding sites are marked as red boxes.
